# Rewilding beyond the wilderness: Beavers can restore stream biodiversity from urban to agricultural to natural landscapes

**DOI:** 10.1101/2025.09.03.674048

**Authors:** Valentin Moser, Silvan Minnig, Leonardo Capitani, Steffen Boch, Nina Cramer, Oriana Edman, Patrick Hofmann, Alex Hürbin, Martin K. Obrist, Christopher Robinson, Dominic Tinner, Luca Zehnder, Christof Angst, Francesco Pomati, Anita C. Risch

**Affiliations:** Community Ecology, Swiss Federal Institute for Forest, Snow and Landscape Research WSL, Birmensdorf, Switzerland; Department of Aquatic Ecology, Eawag: Swiss Federal Institute of Aquatic Science and Technology, Dübendorf, Switzerland; umweltbildner.ch, Bern, Switzerland; Biodiversity and Conservation Biology, Swiss Federal Institute for Forest, Snow and Landscape Research WSL, Birmensdorf, Switzerland; info fauna – Nationale Biberfachstelle, Neuchâtel, Switzerland

**Keywords:** urban, agricultural, Castor, aquatic-terrestrial, rewilding, floodplain, wetlands, ecosystem engineer, restoration

## Abstract

1 Beavers have been promoted as a cost-effective nature-based solution to restore degraded stream ecosystems and enhance overall biodiversity. However, the extent to which beaver-engineering boosts biodiversity in human-impacted landscapes remains poorly understood.
2 We assessed the responses of aquatic and terrestrial taxa (amphibians, dragonflies, fish, macrophytes, plankton, terrestrial vegetation, bats, aquatic, flying and terrestrial invertebrates) to beaver-engineering along a gradient of human land-use intensity in Switzerland. In 16 streams, we measured species richness, abundance, unique species richness, evenness, beta and gamma diversity in a beaver-engineered site with beaver dams and a control site without beaver activity.
3 Especially aquatic and semi-aquatic species profited from beaver-engineering. In response to beaver-engineering, amphibians, dragonflies, macrophytes, and plankton increased in species richness, abundance and unique species richness, while fish and terrestrial plants increased in species richness and unique species richness. Bats only increased in unique species richness, while for the other terrestrial taxa, we found no difference between beaver-engineered and control sites. Community evenness remained largely unaffected across all groups.
4 Overall, beaver-engineering increased biodiversity across a land-use gradient. For macrophytes and amphibians (species richness, unique species richness and abundance), as well for dragonflies (species richness and unique species richness), increases were smaller under higher land-use intensity. For the remaining taxa and metrics, we found no relationship between land-use intensity and responses to beaver engineering.
5 However, for beta-diversity, we found that sites with a high land-use intensity had a lower component of nestedness and stronger community turnover in comparison to more natural sites, with a significant effect for macrophytes. Gamma species richness was generally unaffected by land-use intensity, except for amphibians, which were often only found in beaver-engineered sites in high- land-use intensity areas.
6 *Synthesis and applications*. Our findings demonstrate that beaver-engineering significantly enhances local biodiversity across aquatic and terrestrial ecosystems, even at sites with high land-use intensity. Hence, beavers can effectively restore stream biodiversity from urban to agricultural to natural landscapes. Integrating beaver-engineering into river restoration strategies can substantially advance biodiversity goals under frameworks such as the EU Water Framework Directive and national conservation policies.

## 1 INTRODUCTION

Habitat restoration is a key strategy for reversing biodiversity declines in ecosystems (Díaz et al., 2019). Aquatic ecosystems are particularly affected by habitat loss and degradation (IPBES, 2019) but are challenging to restore due to their complex hydrological dynamics, reliance on upstream conditions, and sensitivity to pollution and climate change (Haase et al., 2025; Palmer et al., 2005). Restoration is further constrained by public opinion, competing land-use interests, and insufficient policy and financial support (IPBES, 2019; Menz et al., 2013). To be effective on a large scale, restoration measures must be cheap, scalable, and socially accepted (Palmer et al., 2005). Rewilding of charismatic species offers a promising solution for restoring ecosystems, balancing ecological benefits with societal acceptance and low long-term management costs (Byers et al., 2006; Du Toit & Pettorelli, 2019; Mutillod et al., 2024; Perino et al., 2019). Among these charismatic species, ecosystem engineers such as the beaver (*Castor fiber* in Eurasia and *C. canadensis* native in North America) are particularly effective in restoring degraded ecosystems (Brazier et al., 2021; Law et al., 2019).

Rewilding with beavers is increasingly recognized as a nature-based solution to enhance biodiversity and ecosystem functioning (Veríssimo & Roseta-Palma, 2023). Physical modifications of the landscape by beavers support both aquatic and terrestrial biological communities by creating habitat heterogeneity and productivity (Rosell et al., 2005). Beaver dams convert lotic (flowing) streams into lentic (standing or slowly flowing water) ponds, expanding slow-water areas by creating riparian zones and depositional environments (zones that accumulate fine sediments) (Law et al., 2016). The ponds accumulate organic matter, supporting large patches of aquatic vegetation and a greater richness of plankton (Czerniawski & Kowalska-Góralska, 2018; Willby et al., 2018). Typically, these heterogeneous aquatic habitats are then populated by aquatic or semi-aquatic taxa like fish, dragonflies and amphibians that show higher richness and abundance compared to non-beaver engineered systems (Larsen et al., 2021; Sommer et al., 2019). Aquatic invertebrates have a more heterogeneous response to beaver-engineering characterised by a high species turnover, but not necessarily a higher species richness (Washko et al., 2022). For terrestrial species, beavers- engineered flooding increases standing deadwood, which leads to a more open canopy (Francomano et al., 2021; Misiukiewicz et al., 2018). Plant richness increases, and flooded sites slowly develop into wetlands (Willby et al., 2018). Bats benefit from beaver-engineering as well (Nummi et al., 2011), while the response of terrestrial arthropods is highly dependent on the taxonomic group investigated (Andersen et al., 2024; Orazi et al., 2022). However, comparing aquatic and terrestrial responses is difficult, as few studies (but see Orazi et al., 2022) have assessed multiple taxonomic groups in beaver-engineered compared to control sites simultaneously.

While the overall increase in local species richness due to beaver-engineering is well studied, the mechanisms behind this community change have only been partially explored. Existing research indicates that beaver-engineering raises landscape-scale (gamma) species richness in communities such as aquatic invertebrates and terrestrial plants by creating new habitats and ecological niches (Czerniawski & Kowalska-Góralska, 2018; Washko et al., 2022; Willby et al., 2018). For beta-diversity, turnover seems to be the main driver, with higher turnover rates both within (Willby et al., 2018) and between beaver-engineered ecosystems compared to sites without beaver-engineering (Law et al., 2019; Wright et al., 2002). Also, human influence can be a strong driver in turnover dynamics, as land-use change can alter species pools and habitat availability (Socolar et al., 2016). However, how human influence and beaver-engineering interact to shape turnover has so far not been assessed.

One potential environmental driver is human land-use, a major cause of habitat and biodiversity loss (IPBES, 2019). Beavers are increasingly colonizing such landscapes with high human land-use, including agricultural and urban areas (Angst et al., 2023; Bailey et al., 2019; Campbell-Palmer et al., 2021; Halley et al., 2020). However, it remains unclear whether beaver-engineering can also increase biodiversity in these highly human-dominated areas. Most studies on beavers’ influences on biodiversity were conducted in natural ecosystems (Washko et al., 2022), where human land-use is minimal. Initial findings from heavily modified agricultural streams suggest that beavers can increase biodiversity even under such conditions (Brazier et al., 2021; Law et al., 2016; Robinson et al., 2020). Even less research was done in urban streams, where beavers create wetland habitats beneficial for people and wildlife (Ciach et al., 2023; Kleinschroth, 2022), but biodiversity gains could be even more constrained by pollution and structural barriers (Kemp et al., 2012). Assessing to what extent beaver-engineering impacts biodiversity in human-dominated landscapes is therefore essential for understanding the full potential of beavers in restoring degraded ecosystems. As beavers continue to expand into areas with high land-use intensity, such knowledge can inform conservation policy, land-use planning, and nature-based restoration strategies.

In this study, we assessed to what extent beaver-engineering (dam building) affects plant and animal species richness, abundance, unique species richness, evenness, nestedness and gamma species richness across 16 stream ecosystems that span a gradient of land-use intensity. We collected data for both aquatic and terrestrial taxa in a beaver-engineered area (pool) and an area without beaver-engineering (control) at each stream site. First, we hypothesised (H1) that beaver-engineering leads to overall higher richness, abundance, unique species richness of the different taxa, and to more equitable communities (higher evenness) compared to control, reflecting the variety of niches and resources introduced by beaver-engineering (Rosell et al., 2005). Second, we hypothesised (H2) that biodiversity gains from beaver-engineering would be most pronounced at sites with higher human land-use intensity due to their low initial habitat complexity (Law et al., 2016). Third, we hypothesised (H3) that species nestedness is lower in natural systems, as high human land-use increases nestedness (Wang & Han, 2023). Fourth, we hypothesised (H4) that gamma species richness is highest in natural systems due to a lower total species pool in high land-use areas (Pärtel et al., 2025).

## 2 MATERIALS AND METHODS

### 2.1 Study area

This study was conducted in 16 stream ecosystems distributed across the Swiss midlands (Fig. S1, Table S1). The selected sites encompassed a range of typical beaver-engineered habitats with dams in Switzerland, spanning higher to lower order streams and situated in urban, agricultural, and natural landscapes (with a gradient in land-use intensity). All sites had a minimum of four years (with a maximum of 14 years) of beaver-engineering to allow sufficient time for the beaver-engineering effects to manifest.

At each site, we established a paired design: A beaver-engineered area with an active beaver dam (Pool) and a control area (Control) without beaver-engineering located approximately 500 m upstream or downstream of the Pool along the same stream (Fig. S2). The Pool area extended 25 meters downstream (outflow) and 75 meters upstream of the beaver dam (pool), the Control area was also a 100-meter stretch. We chose a difference of 500 m between the Pool and Control to minimize landscape-scale effects (same potential species pool present). We matched the land-use intensity and impacts of humans at Pool and Control of each site as closely as possible to ensure comparability. For example, in regulated streams, both Pool and Control included artificial streambeds. In the first year of the study, some unusually high rainfalls led to the destruction of 5 of 8 dams. Nevertheless, as beaver-engineering effects were still evident, we continued with data collection.

### 2.2 Data collection

We collected data for 10 different aquatic and terrestrial plant and animal taxa within the 16 stream ecosystems over two years, with eight sites sampled in 2021 and eight in 2022. Sampling methods were designed to account for the different taxa included. Sampling for the fish, macrophytes, amphibians, dragonflies (including damselflies) and aquatic invertebrates were conducted by ecological consultancies coordinated by the national beaver office (https://www.infofauna.ch/de/nationale-koordinationsstellen/biber) as part of a monitoring project on the effects of beavers on aquatic biodiversity. Plankton, terrestrial flying arthropods, bats, terrestrial invertebrates and terrestrial plants were sampled by the project team of WSL and EAWAG in June and July of the respective year. Flying arthropods and bats were sampled in eight systems in the second year only; all other taxa were sampled at all 16 sites.

Fish were sampled using electrofishing, with at least two rounds per site conducted according to standard Swiss protocols to ensure comprehensive species detection around September (Schager & Armin, 2004). Where a quantitative survey was not possible (4 of 16 sites) due to the depth of the pool, electrofishing was done quantitatively in 30 random locations to assess richness (Vonlanthen et al., 2022). One site (Nider) was done in 2022 instead of 2021. Macrophytes, dragonflies (including damselflies; collectively called dragonflies hereafter), and amphibians were surveyed along the 100-meter sections in both the Pool and Control. Macrophytes were identified to species level with their cover recorded in July or August (Känel et al., 2017). For dragonflies, adults and exuviae were counted during four field surveys conducted between May and August.

Amphibians, including larval stages and egg masses, were identified and counted inside and outside of the water. These surveys were conducted in March during the day, as well as April, May and June during the night. Aquatic invertebrates were sampled using kick-netting, targeting two organic (e.g. macrophytes, deadwood) and two inorganic substrates (e.g. sand, gravel) habitats within the 100 m-long Pool and Control stretch in May, June or July (BAFU (Hrsg.), 2019). All aquatic invertebrates were counted and identified to species level or else to the lowest taxonomic level possible. Plankton was identified as operational taxonomic units (see Table S3 for overview), based on pictures taken with an automated dark-field imaging microscope (Merkli et al., 2024). For this purpose, we passed 50 litres of water collected from the centre of the Pool and Control through the microscope. The pictures were manually annotated and classified (Chen et al., 2025; Merz et al., 2021). Flying arthropods were sampled using flight interception traps (Gossner et al., 2014), constructed with crossed fabrics, measuring 50 cm × 50 cm, with a funnel and collection containers placed at the top and bottom containing 96 % ethanol. They were mounted as low above the water as possible in the middle of the respective stream body (Pool and Control), with traps deployed for 9 days during June and July 2022, each. Samples were stored in ethanol for later processing. Species were counted and identified to order level with the aid of a binocular and literature (Chinery & Falk, 2012). Bats were monitored acoustically for 5 days each in June and July 2022 using automatic recorders placed at the water edge in the same area and at the same time the flying arthropods were sampled. Bat acoustic data was classified using BatScope with a standard protocol (Obrist & Boesch, 2018) to obtain species IDs and abundances. Terrestrial invertebrates and terrestrial plants were sampled in a 5 x 1 m plot located one meter from the water’s edge in the centre of the Pool and Control area. Within each plot, we sampled the invertebrates at the two ends of the 5 x 1 m plot in cylindrical baskets (50 cm diameter, 67 cm height, woven fabric) using suction sampling (Resch et al., 2021) on a sunny day between 10:00-17:00 during peak arthropod activity. Again, the samples were stored in ethanol, individuals were counted and identified to order level with the help of a binocular. Plants were identified to species level, and the cover of each species was recorded within the 5 x 1 m plots. Plant abundance was calculated based on the highest individual cover per species on either herbs, shrubs, or tree cover level.

All relevant cantonal authorities, landowners and governmental agencies were contacted by telephone to obtain permission for access and sampling. A written permission was only necessary for the Talent and Biber site, part of a nature reserve. The fish monitoring was carried out by Ecqua (https://www.ecqua.ch) after obtaining all necessary permissions from the cantonal hunting and fisheries authorities. No additional animal ethic approval was necessary for the work conducted.

### 2.3 Land-use intensity

Land-use intensity data for each study site were extracted from Geodienste.ch and swisstopo.admin.ch using ARCGIS (Pro v. 2.8, Esri Inc., 2021 und QGIS 3.40 ’Bratislava’ 2024) (Table S2). A radius of 250 meters was selected around the centre of each Pool and Control to avoid overlap between paired sampling areas. A visual inspection of the data showed highly reliable classification for agricultural (e.g., crop fields, pastures, areas that farmers maintain for biodiversity) and natural (e.g., forests, riparian areas) land-use. Areas without classified land-use were very often urban areas and human infrastructure, such as streets, railways, and housing. Therefore, missing data was assigned to urban land-use.

### 2.4 Statistical Analysis

We calculated species richness, abundance, unique species richness, and evenness for each of our taxa described above. As the sampling effort was designed to assure equal efforts, we did not rarify the data (Underwood, 1996). Unique species richness is the number species not shared between Control and Pool per site. We also quantified cumulative unique species, which are species found only in Pool or Control over the whole 16 sites. For the aquatic invertebrates, we first redistributed counts from higher-rank identifications (order or higher→ family → genus) proportionally to their lower-rank descendants within each sample, then calculated richness, abundance and diversity indices on the pruned taxa list, and derived unique-taxon counts by comparing the pruned Control and Pool samples at each site while treating ancestor–descendant pairs as overlaps. We assessed the influence of beaver-engineering on the different biodiversity and abundance measures for each taxon with generalized linear mixed-effects models (GLMMs), with site included as a random intercept to account for the paired design and possible site-level heterogeneity. For count data (species richness, abundance, unique species richness), we initially evaluated poisson or negative binomial models using lme4 version 1.1-35.5 (Bates et al., 2015). We adjusted the models, guided by model diagnostics dispersion, zero-inflation, and residual uniformity of the Dharma package version 0.4.6 (Hartig, 2017), as well as lower Akaike’s Information Criterion (AIC), with differences >2 considered meaningful (Table S5). We additionally calculated marginal and conditional R² values as measures of explanatory power. These R² values helped to assess whether improvements in model fit according to AIC translated into meaningful increases in variance explained by the fixed (marginal R²) or fixed and random effects combined (conditional R²). Generally, we used the poisson distribution for species richness and unique species richness and the negative binomial distributions for the modelling of the abundance data. The macrophyte abundance data showed strong overdispersion and zero-inflation and was therefore modelled using a zero-inflated negative binomial GLMM (ZINB) fitted via glmmTMB version 1.1.10 (Brooks et al., 2017). Dispersion tests indicated underdispersion in some poisson models for species richness and unique species richness. However, using negative binomial models did not improve model fit based on AIC or residual diagnostics. Thus, we retained the poisson GLMMs in these cases for parsimony and interpretability. Some models produced singular fit warnings (near-zero variance estimates for random intercepts), but the random structure was maintained to respect the paired design. Evenness, bounded between 0 and 1, was modelled using beta regression with a logit link, again with site as a random intercept. To accommodate boundary values, evenness values were transformed via a squeezing approach (Smithson & Verkuilen, 2006). Zero-inflated beta models were tested, where DHARMa residuals showed strong deviations, but as DHARMa residuals for these models also indicated significant deviations, we retained standard and simpler beta models for all taxa for consistency.

To assess whether beaver-engineering effects on species richness, abundance, unique species richness, and evenness varied along land-use gradients, we calculated the difference in species richness, abundance, unique species richness and evenness between Pool and Control sites (Δ = Pool – Control) per taxa and site. We averaged the sum of agricultural, urban, and natural land-use cover types per site across Pool and Control. Agricultural and urban land were summarized into an overall human impact called land-use intensity. We fitted separate linear models for each biodiversity metric (and land-use type (agriculture, urban, natural, and overall land-use intensity), independently for each of our 10 taxa. For each model, we assessed whether land-use intensity significantly predicted changes in biodiversity measures between Pool and Control. For visualization, the land-use variables were normalized [–1, 1] to allow for comparison across taxa.

To assess whether beaver-engineering effects on gamma species richness and beta diversity varied along land-use gradients, we quantified total gamma species richness per site and calculated pairwise beta diversity between Pool and Control for each site and taxa using the Sørensen index, partitioned into turnover and nestedness. For the statistical evaluations of tunrover and nestedness, we use the nestedness component, which equals nestedness divided by total beta-diversity. We also fitted one model per diversity index and taxa. We fitted simple linear models to test for direct relationships between biodiversity and diversity metrics and land-use predictors. We deliberately chose parsimonious models to answer our hypotheses, acknowledging that these represent simplified representations of complex ecological systems. All models and visualizations were conducted in R v4.4.2 using ggplot2 version 3.5.1 (Wickham, 2016). To check code and improve the flow and grammar of the writing, ChatGPT (model 4.o) was used.

## 3 RESULTS

First, we analysed richness, abundance, unique species richness and evenness (Fig. 1, Table S4). Species richness and unique species richness per site were significantly higher for amphibians, dragonflies, fish, macrophytes, plankton, and terrestrial plants in beaver-engineered Pool compared to Control areas (p < 0.05, Fig. 1, Table S4). Additionally, unique species richness was higher in the Pool compared to the Control for bats (p < 0.01, Fig. 1, Table S4). No significant differences were detected between Pool and Control for aquatic invertebrates, bats, terrestrial invertebrates, and flying arthropods (Fig. 1, Table S4). Abundances were higher in Pool compared to Control for macrophytes (p < 0.05), dragonflies (p < 0.01), plankton (p < 0.05), and amphibians (p < 0.001). For evenness, only flying arthropods (p < 0.05) showed a statistical difference between the Pool and Control.

**Fig. 1.**
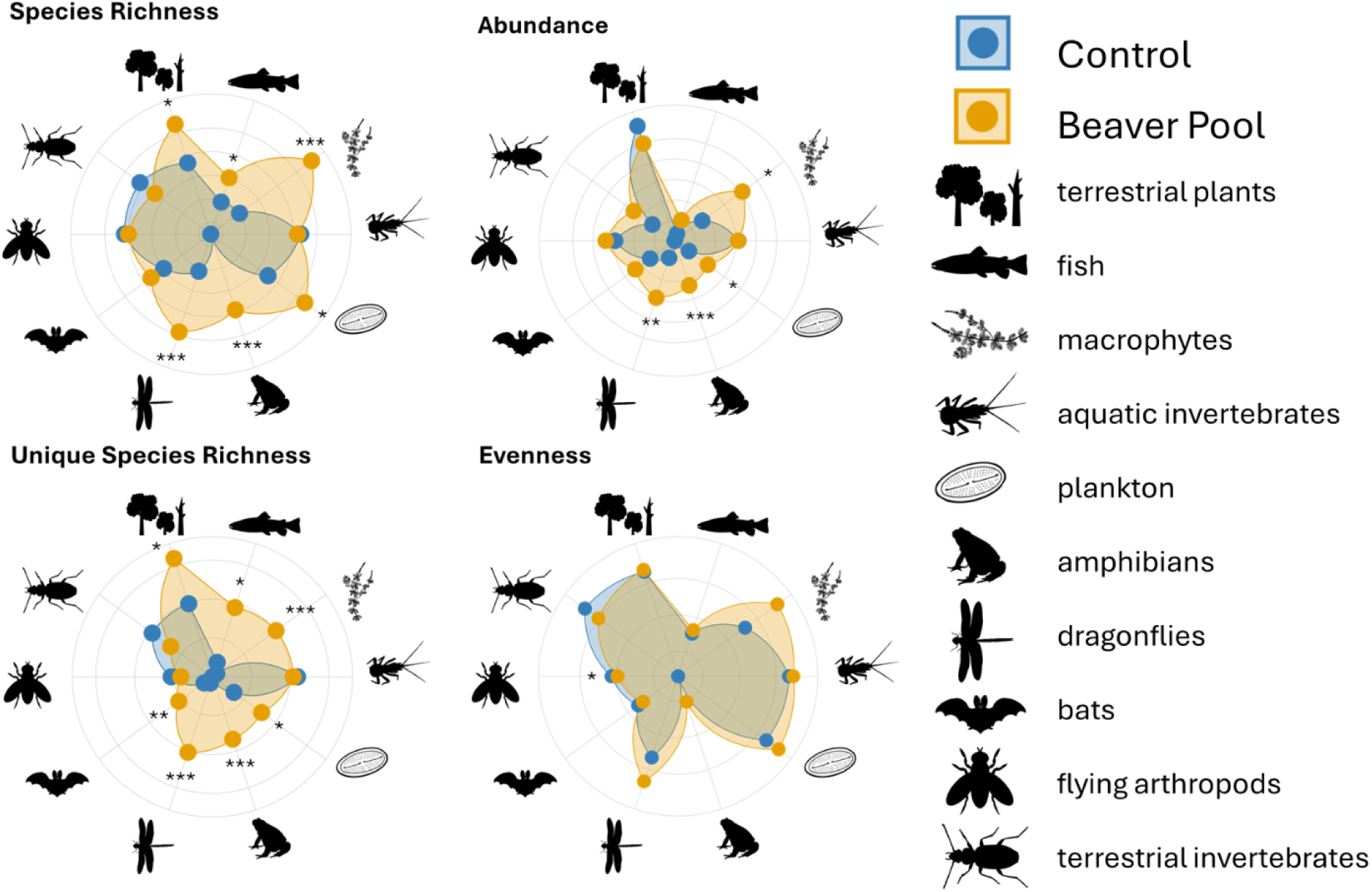
Radar plots showing the differences in species richness, abundance, unique species richness, and evenness between paired Pool (orange, with beaver-engineering) and Control (blue) areas. All values were normalized. Significance differences between Pool and Control are indicated with stars: *= p < .05, ** = p < .01, *** = p < .001. Flying arthropods and bats were sampled in one year only (n = 8), while all other taxa were sampled in both years (n = 16).

Secondly, we tested the interaction between human land-use intensity and beaver-engineering on biodiversity (Fig. 2). Amphibian, dragonfly and macrophyte richness and unique species richness decreased significantly with higher land-use intensity (Fig. 2, Table 1). As Δ richness and Δ unique species richness are the same, we do not show the results for Δ unique species richness. Abundance was less affected by land-use intensity than species richness, but we found significantly lower Δ abundance of amphibians and macrophytes in Pool compared to Control areas with higher land-use intensity (Fig. 2, Table 1). Some models did not perform well, showing the following assumption violations: ΔFish abundance failed the uniformity test, while all taxa except dragonflies exhibited zero-inflation for Δ richness and Δ unique species richness. Zero-inflation was also detected for Δ abundance and Δ evenness of amphibians, fish and plankton. As outlined in the models, more complex models did not improve model fit and therefore the more parsimonious models were retained.

**Fig. 2.**
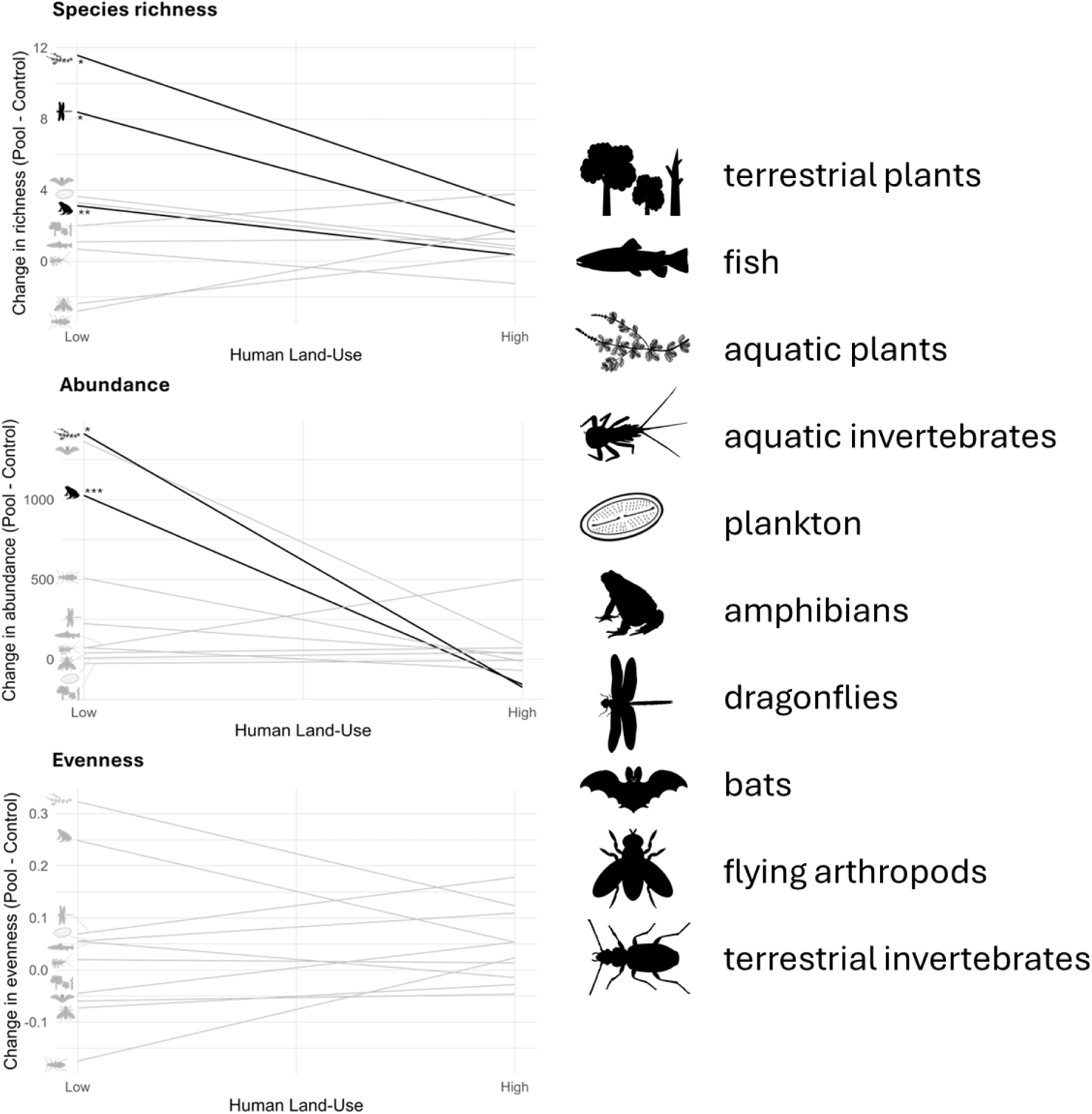
Linear regressions for Δ (Pool - Control) species richness, abundance and evenness for each taxa in response to overall land-use intensity (agricultural and urban land-use combined). Regression lines in bold represent significant taxa, the levels of significance are indicated with stars: * = p < .05, ** = p < .01, *** = p < .001. Values above the zero line stand for a positive effect of beaver-engineering. Δ unique species richness is the same as Δ species richness. For this reason, model results are only shown for Δ species richness.

**Table 1:** Linear model results for Δ (Pool - Control) species richness, abundance and evenness for each taxa in response to overall land-use intensity. R² and adj. R² = model fit indicators, and p-value = significance of the relationship. Land-use intensity is the sum of urban and agricultural land cover. Δ unique species richness is the same as Δ species richness. For this reason, model results are only shown for Δ species richness.

| **Metric** | **Community** | **Estimate** | **Std.error** | **R.squared** | **Adj.r.squared** | **p.value** |
| --- | --- | --- | --- | --- | --- | --- |
| richness | amphibians | -1.44E-05 | 4.15E-06 | 0.46 | 0.42 | **<0.01** |
| richness | aquatic invertebrates | -1.01E-05 | 2.28E-05 | 0.01 | -0.06 | 0.66 |
| richness | flying arthropods | 2.41E-05 | 1.04E-05 | 0.47 | 0.38 | 0.06 |
| richness | terrestrial invertebrates | 1.44E-05 | 1.37E-05 | 0.07 | 0.01 | 0.31 |
| richness | bats | -1.47E-05 | 1.41E-05 | 0.15 | 0.01 | 0.34 |
| richness | dragonflies | -3.53E-05 | 1.21E-05 | 0.38 | 0.33 | **<0.05** |
| richness | fish | 8.45E-07 | 5.15E-06 | 0.00 | -0.07 | 0.87 |
| richness | macrophytes | -4.41E-05 | 1.77E-05 | 0.31 | 0.26 | **<0.05** |
| richness | plankton | -1.37E-05 | 1.23E-05 | 0.08 | 0.02 | 0.28 |
| richness | terrestrial plants | 9.37E-06 | 1.17E-05 | 0.04 | -0.02 | 0.44 |
| abundance | amphibians | -6.19E-03 | 1.45E-03 | 0.57 | 0.54 | **<0.001** |
| abundance | aquatic invertebrates | -7.46E-04 | 1.00E-03 | 0.04 | -0.03 | 0.47 |
| abundance | flying arthropods | 1.66E-04 | 6.02E-04 | 0.01 | -0.15 | 0.79 |
| abundance | terrestrial invertebrates | -2.73E-03 | 2.86E-03 | 0.06 | -0.01 | 0.36 |
| abundance | bats | -6.62E-03 | 7.26E-03 | 0.12 | -0.02 | 0.40 |
| abundance | dragonflies | -1.00E-03 | 7.17E-04 | 0.12 | 0.06 | 0.18 |
| abundance | fish | 2.25E-03 | 4.22E-03 | 0.02 | -0.05 | 0.60 |
| abundance | macrophytes | -8.30E-03 | 3.55E-03 | 0.28 | 0.23 | **<0.05** |
| abundance | plankton | 1.89E-04 | 2.73E-04 | 0.03 | -0.04 | 0.50 |
| abundance | terrestrial plants | 1.05E-04 | 1.82E-04 | 0.02 | -0.05 | 0.58 |
| evenness | amphibians | -1.02E-06 | 1.56E-06 | 0.03 | -0.04 | 0.52 |
| evenness | aquatic invertebrates | -3.21E-08 | 8.65E-07 | 0.00 | -0.07 | 0.97 |
| evenness | flying arthropods | 2.35E-07 | 4.04E-07 | 0.05 | -0.10 | 0.58 |
| evenness | terrestrial invertebrates | 1.04E-06 | 1.11E-06 | 0.06 | -0.01 | 0.36 |
| evenness | bats | 6.92E-08 | 8.27E-07 | 0.00 | -0.17 | 0.94 |
| evenness | dragonflies | 5.70E-07 | 1.74E-06 | 0.01 | -0.06 | 0.75 |
| evenness | fish | -3.65E-07 | 9.23E-07 | 0.01 | -0.06 | 0.70 |
| evenness | macrophytes | -1.05E-06 | 1.78E-06 | 0.02 | -0.05 | 0.57 |
| evenness | plankton | 2.83E-07 | 1.84E-06 | 0.00 | -0.07 | 0.88 |
| evenness | terrestrial plants | 5.14E-07 | 7.82E-07 | 0.03 | -0.04 | 0.52 |

Thirdly, we assessed how natural, agricultural, and urban land use separately influence biodiversity in beaver-engineered streams (Fig. S3, S4). With higher agricultural land-use, Δ richness of amphibians and dragonflies was negative, while higher natural land cover had a positive effect on Δ richness of amphibians, dragonflies and macrophytes (Fig. S3). With higher urban land-use, Δ richness and unique species richness increased for terrestrial invertebrates (Fig. S3, Table S6). High agricultural land cover had a negative effect for or Δ abundance of amphibians, while high natural land cover was positive for Δ abundance of amphibians and macrophytes. For Δ evenness, no significant effects of agricultural, urban or natural land-use were detected for any of our taxa (Fig. 2, Fig. S3).

Lastly, to understand the mechanisms behind the change in species richness across the 10 taxa, we assessed cumulative unique species, beta diversity (ratio of nestedness and turnover) and gamma species richness separately for each taxa. We recorded 148 cumulative unique species in Pool, compared to 79 in Control (Table S7). This cumulative count includes the highest resolved taxonomic levels, mostly species, but occasionally higher taxonomic levels such as genera or orders. The highest numbers of cumulative unique species were found for aquatic invertebrates (34 Pool, 29 Control) and terrestrial plants (55 Pool, 37 Control). Other taxa contributed fewer cumulative unique species: Macrophytes (19 Pool), dragonflies (14 Pool), plankton (12 Pool, 4 Control), fish (7 Pool, 1 Control), terrestrial invertebrates (2 Pool, 3 Control), amphibians (2 Pool), flying arthropods (2 Control), and bats (1 Pool). For the nestedness component, amphibians could not be evaluated due to a low sample size especially in control areas. Seven of the remaining nine taxa showed a negative relationship with land-use intensity, but only the relationship between land-use intensity and macrophytes was significant (R² = 0.52, p < 0.01, Table S8). For gamma species richness, amphibians showed a significantly smaller gamma species richness with increasing land-use intensity (R² = 0.53, p < 0.01, Table S8).

## 4 DISCUSSION

Here, we presented results from a large-scale field study testing the influence of beaver-engineering on the biodiversity of different aquatic and terrestrial taxa across a land-use gradient. Our results showed that beaver-engineering enhanced biodiversity and species abundances, particularly for aquatic taxa, across a gradient of land-use. We found significant increases in species richness, abundance and unique species richness in beaver-engineered areas compared to nearby controls.

However, we did not find consistent increases in evenness across taxa, indicating that while beaver- engineering increased species numbers, it did not alter species dominance patterns (Fig. 1). Increases in species richness, abundance and unique species richness extended to high land-use intensity systems, but were often weaker than in more natural systems, suggesting that the benefits of beaver-engineering for restoration purposes may be constrained by the lower habitat quality and depleted species pools typically found under high human land-use. Lastly, with increased land-use, the component of nestedness of the total beta diversity decreased (Fig. 3). Overall, our results highlight the ecological role of beaver-engineering in promoting multi-taxa biodiversity across aquatic and terrestrial taxa, even in areas with high land-use intensity.

**Fig. 3.**
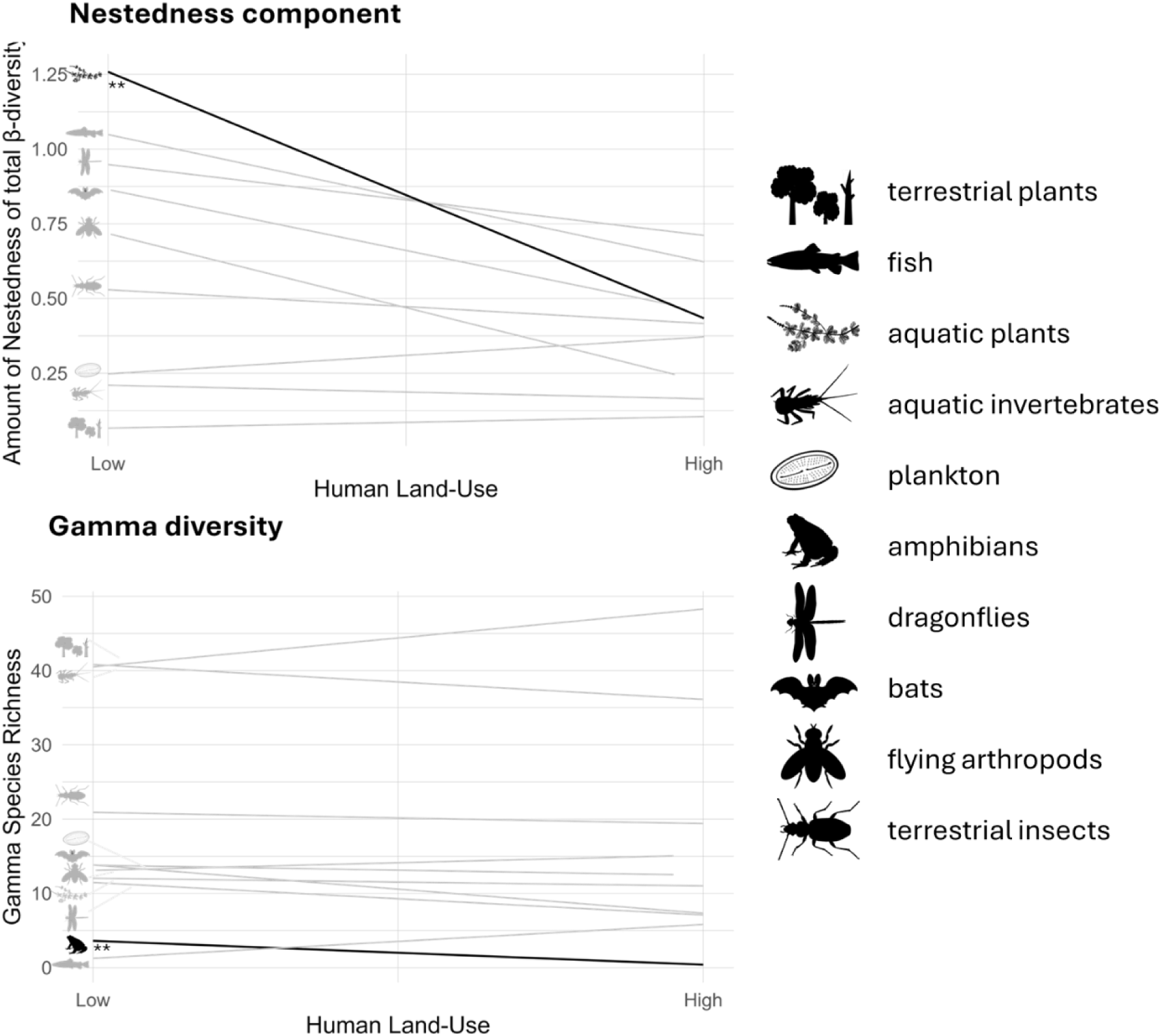
Linear regression with the component of nestedness of the total beta diversity between Pool and Control, and total (gamma) species richness per site in response to overall land-use intensity (agricultural and urban land-use combined). Regression lines in bold show taxa for which we found significant relationships with land-use intensity, while the strength of the relationship is indicated with stars * = p < .05, ** = p < .01, *** = p < .001. For beta diversity, higher values indicate a stronger role of nestedness relative to turnover. For gamma species richness, values represent the total species pool (Pool + Control) per site. Significant effects of land-use intensity are indicated with stars. Due to the absence of amphibians in most Control sites, we excluded this group from the nestedness component analyses.

### 4.1 Responses of aquatic and terrestrial taxa to beaver-engineering

In our study, we recorded many cumulative unique species only in the beaver-engineered Pool (Table S7), most notably several species each for amphibians, macrophytes, and dragonflies, which reinforces previous observations that beaver-engineering creates or re-establishes habitat niches for species that are otherwise unable to inhabit uniform stream channels (Romansic et al., 2021).

However, in our Control areas we also found some cumulative unique species, most numerous for aquatic invertebrates and terrestrial plants. This supports earlier findings that beaver-engineering can displace certain types of habitats and their associated species, altering community composition, but not necessarily increasing local alpha richness (Washko et al., 2022). Flying arthropods had lower evenness in the Pool compared to Control areas, possibly due to the high abundance of aquatic insects like Chironomidae that can emerge from beaver pools (Nummi et al., 2011). Evenness remained unaltered by beaver-engineering, but for flying arthropods. This suggests that species gains were driven by the addition of species rather than increases in the dominance of a few species, supporting the idea of beavers as keystone ecosystem engineers (Yeakel et al., 2020).

Our results show that beavers generally increase species richness, abundance and unique species richness across different taxa, supporting previous findings. However, with our design of sampling aquatic and terrestrial taxa simultaneously, we were also able to compare differences in responses of aquatic and terrestrial taxa to beaver-engineering. We found significant increases in species richness, abundance and unique species richness for most aquatic species with beaver- engineering. The exceptions were aquatic invertebrates, for which we found mixed results. This is similar to what is reported in literature (Brazier et al., 2021; Law et al., 2016; Orazi et al., 2022; Rosell et al., 2005; Washko et al., 2022). In contrast, for the terrestrial taxa, we only found a higher species richness for plants and a higher unique species richness for bats in our beaver-engineered systems. This is also consistent with earlier findings (Ciechanowski et al., 2011; Law et al., 2016; Orazi et al., 2022). These differences between aquatic and terrestrial taxa may reflect how beavers modify their surroundings in aquatic compared to terrestrial ecosystems. In the water, beaver-engineering substantially alters water area, speed and sediment disposition (Larsen et al., 2021), while on land, changes include higher light availability, increased soil moisture and exposed soil (Law et al., 2016). The more thorough physical transformation of the aquatic habitats may explain the stronger and more consistent response of aquatic taxa to beaver-engineering. On land, species gains and losses may balance out: Some terrestrial ground-dwelling arthropods may be displaced, while others profit from an increase in detritus and a higher plant species richness (Andersen et al., 2024). Our lack of species-level identification for terrestrial invertebrates could obscure such finer patterns. It is also possible that our results reflect the earlier succession stages of the beaver systems we studied (4-14 years old). Later successional stages, such as wet meadows or standing deadwood, may provide greater benefits for terrestrial taxa like butterflies or saproxylic beetles (Andersen et al., 2024).

Notably, the taxa responding most strongly to beaver-engineering in our study, amphibians and dragonflies, are semi-aquatic, highlighting how beaver-engineering influences taxa across aquatic-terrestrial boundaries.

### 4.2 Responses of different taxa to beaver-engineering with increasing land-use intensity

Land-use intensity had a weaker effect on most taxa than we expected. However, this still is good news: We found that beaver-engineering increased richness, abundance and unique species richness even in highly human-modified landscapes. The strongest negative effects of high human land-use were observed for macrophytes, amphibians and dragonflies, taxa known to be particularly sensitive to pollution and land-use changes (Babko et al., 2023; Hopkins, 2007; Samways et al., 2025). This observed negative effect of land-use on these taxa is consistent with the idea that less natural landscapes offer smaller regional species pools and reduced habitat connectivity, limiting colonization of new (wetland) habitats (Law et al., 2016). Physical constraints in high human land-use landscapes may also limit the extent of beaver-engineering on different taxa. For example, an earlier study showed that a narrow, incised agricultural stream showed smaller increases in habitat area and species diversity than a broad-valley stream where beavers created a large wetland complex (Robinson et al., 2020). Similarly, in confined, artificial riverbeds, we observed that beaver dams were flushed out more easily by extreme weather events, limiting the long-term stability of beaver-engineered ecosystems. Nevertheless, even in quite degraded streams such as some of the ones included in our study (Zeh Weissmann et al., 2009), beaver-engineering still provided substantial benefits for amphibians, macrophytes, and dragonflies, albeit smaller than in more natural habitat.

One other noteworthy result of our study is the increase of terrestrial arthropod species richness in urban areas (Fig. S3, Table S6), indicating that beaver-engineering can provide substantial ecological value even in heavily modified settings for some taxa. While our simple models revealed some significant relationships between land-use intensity and beaver-engineered species richness, abundance and unique species richness, we acknowledge that these models only explain a part of the variance found in our biodiversity measures. Residual patterns suggest that additional factors, such as stream size (Ciach et al., 2023), can influence these relationships. It is further important to note, that even natural systems in Switzerland are shaped by human land-use, such as historic drainage ditch installation, possibly still influencing current responses.

### 4.3 Beaver-engineering affecting beta and gamma species richness

We found a lower nestedness component with increasing land-use intensity for 7 of 9 taxa (amphibians not evaluated due to small sample size), but the result was only statistically significant for the macrophytes. This contrasts with previous results reporting an increasing nestedness compared to turnover of taxa with higher human land-use (Wang & Han, 2023). Human land-use often filters species communities to species that are particularly resilient (Keck et al., 2025; Moi et al., 2023). Beaver-engineering seems to be able to partly reverse that filtering and allow a novel community to form. For gamma species richness, only amphibians were negatively affected by increasing human land-use intensity: In many high land-use intensity control sites we found no amphibians, therefore only the beaver-engineered area contributed to amphibian gamma richness. Amphibians are particularly sensitive to high land-use intensity (Hopkins, 2007), but beaver ponds also offer preferred slower-flowing water areas rather than streams (Schmidt et al., 2023). Overall, our beta- and gamma-diversity results highlight the potential of beaver-engineering to increase species diversity even in landscapes with high human land-use intensity: In other words, beavers create the conditions for some species to establish where they otherwise might not.

### 4.4 Application

We demonstrated that beaver-engineering enhances species richness, abundance and unique species richness, particularly for aquatic taxa, making beavers a valuable partner and highly effective tool for conservation and restoration. By creating and maintaining diverse habitats across aquatic– terrestrial boundaries, they act as true ecosystem engineers (Jones et al., 1994). Our findings add to the growing recognition that rewilding ecosystem engineers can complement or even substitute costly mechanical restoration measures. As scalable and self-sustaining contributions to freshwater restoration, beavers represent a promising nature-based solution for counteracting biodiversity loss.

Importantly, taxon gains from beaver-engineering are not limited to natural areas: Even in agricultural and urban landscapes, we found higher species richness and abundances, as well as novel species communities at the landscape scale. In Switzerland, this potential is partially recognized in direct management recommendations in forest biodiversity promotion programs (BAFU, 2023).

More broadly, restoration of streams via beaver-engineering aligns with the goals of the Swiss Water Protection Act (GSchG) and the EU Water Framework Directive, aiming to restore the ecological integrity of freshwater ecosystems. However, the potential of beaver-engineering to support ecosystem restoration is currently often constrained by human-wildlife conflicts. Without sufficient riparian buffers, beaver-engineering can lead to flooded fields, roads, buildings or gardens (Angst et al., 2023; Pollock et al., 2015). In the Swiss agricultural areas, 75% of streams have a road in direct proximity (Angst, 2010). This limits the terrestrial influences of beavers and often results in management interventions such as dam removals and burrow destruction, which further restricts the potential of beavers to contribute to ecosystem restoration in time and space. Careful, site-specific management is therefore needed to balance between the potential of beaver-engineered nature-based solutions and the prevention of infrastructure damage or agricultural losses.

## ACKNOWLEDGEMENTS

We thank the towns and cities, government agencies, and landowners for granting access and permission to use their land as study sites. We thank the involved ecological consultancies and biologists for their work in the field, particularly Thomas Kreienbühl and his helpers (ecqua.ch), Daniel Küry, Raphael Krieg, Pascal Schweizer und Alexander Freude (lifescience.ch), Thomas Mathis, Beatrice Lüscher, Marco Thoma, Jérôme Pellet, Melanie Annen und Silvia Zumbach (karch.ch), Timon Polli and all other helpers who supported us during the field days. We also thank Julia Holmes, Tobias Marfil, Sérena Hadfield and all other helpers for their contribution during the field season and sorting of invertebrates. This study was funded by the Federal Office for the Environment (FOEN) and the ETH Board as part of the Blue-Green Initiative of ETH Zurich.

## AUTHOR CONTRIBUTIONS

Valentin Moser, Silvan Minnig, Christof Angst, Anita C. Risch and Francesco Pomati conceived the ideas and designed the study; Valentin Moser, Steffen Boch, Nina Cramer, Oriana Edman, Patrick Hofman, Alex Hürbin, Dominic Tinner and Luca Zehnder contributed to the data collection; Martin K. Obrist verified bat echolocation calls; Valentin Moser analysed the data; Valentin Moser led the writing of the manuscript; Anita C. Risch, Francesco Pomati, Christopher Robinson and Martin C. Obrist supervised in the project; Anita C. Risch and Francesco Pomati jointly wrote the proposal for funding, organized and led the overall study. All authors contributed critically to the drafts and gave final approval for publication.

