## Supplementary Figures and Tables for "Rewilding beyond the wilderness: Beavers can restore stream biodiversity from urban to agricultural to natural landscapes"

**SUPPORTING INFORMATION**

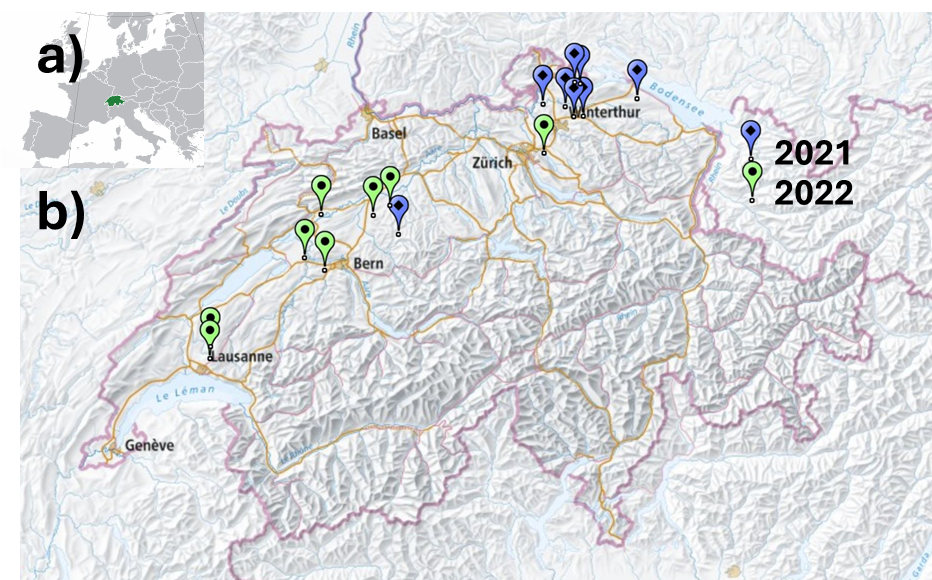

*Fig. S1 Location of the 16 study sites distributed across the Swiss midlands. a) map of Europe with Switzerland shown in green. b) map of Switzerland with the location of the eight sites sampled in 2021 shown with a blue marker and the eight sites sampled in 2022 with a green marker. Background: By Hayden120 and NuclearVacuum - Location European nation states.svg, CC BY-SA 3.0,* [*https://commons.wikimedia.org/w/index.php?curid=8109256*](https://commons.wikimedia.org/w/index.php?curid=8109256) *and ©swisstopo*

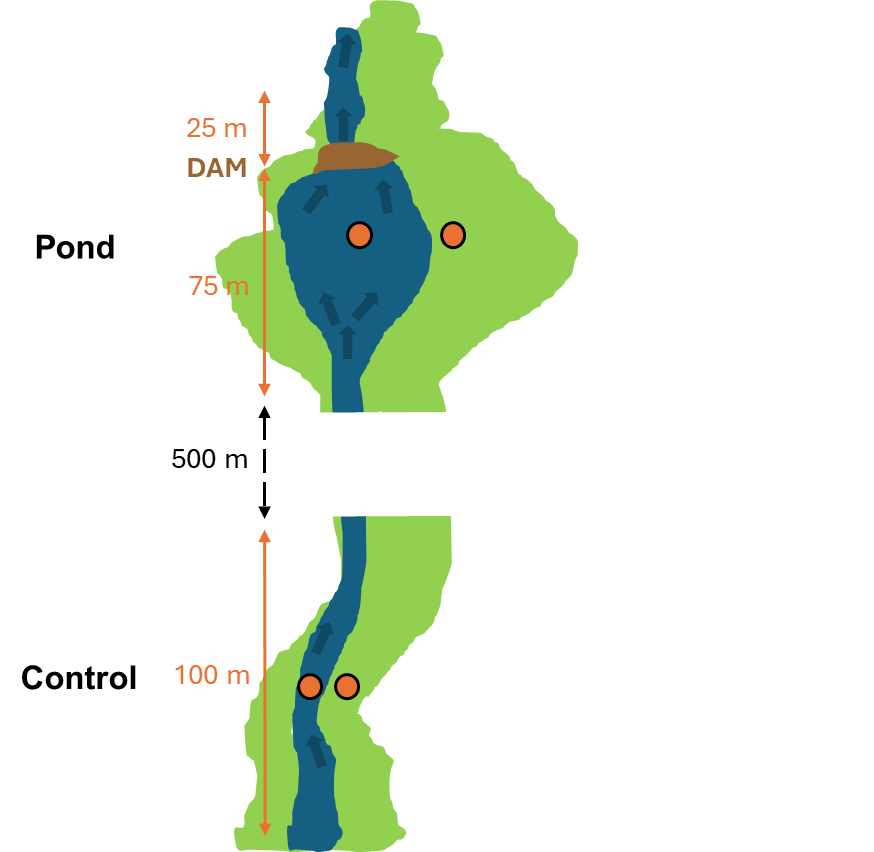

*Fig. S2 Experimental design. Shown are Control and Pool areas along the same stream and the beaver dam (brown). Sampling was conducted along two 100-meter stretches within the water and in the adjacent terrestrial habitat. Sampling locations are in orange: 100 metres two-headed arrows for taxa like fish, single aquatic point for taxa like plankton and single terrestrial points for taxa like bats. For the Pool the 100 meter sampling area extended from 25 meters downstream of the beaver dam to 75 meters above the dam. The Control is located approximately 500 m up- or downstream of the respective Pool (see Table S1).*

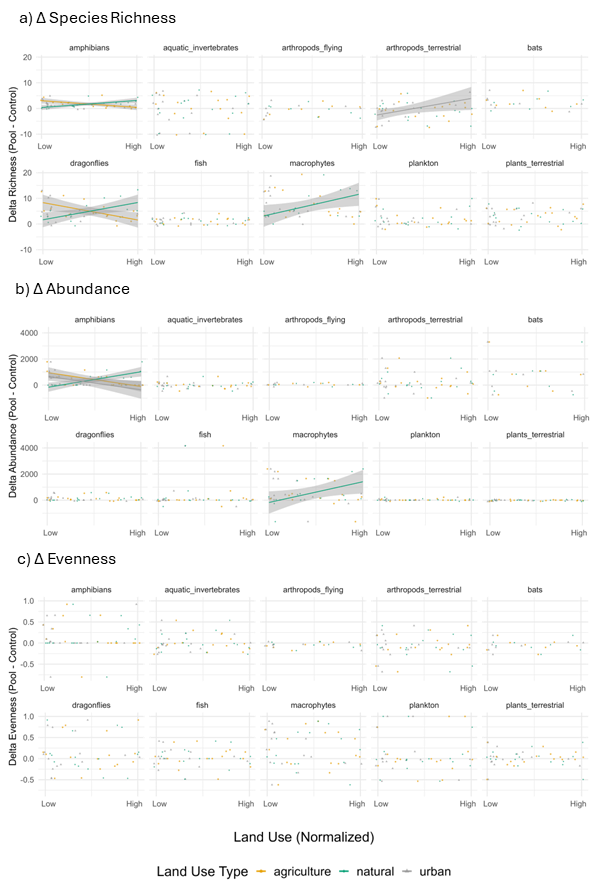

*Fig. S3 The change in* Δ *for a) species richness, b) abundance and c) evenness for each community across sites.* Δ *are shown for three land-use types, agricultural (orange), natural (green) and urban (grey).* Δ *unique species richness is the same as richness and therefore not shown separately. In the main results and text, agriculture and urban were summarized to “land-use intensity”.*

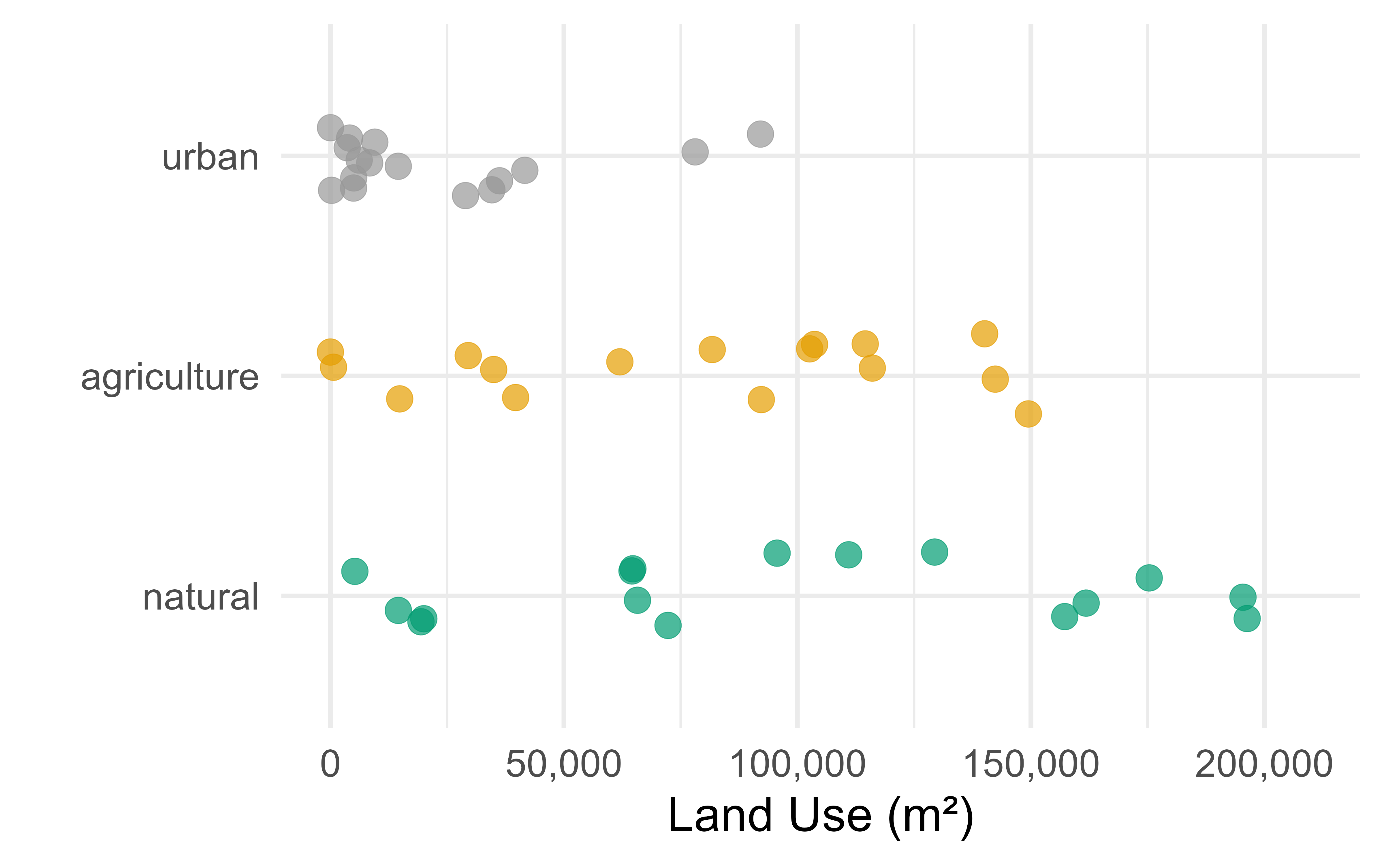

*Fig. S4 Distribution of land-use intensity across the study sites. Each point represents the averaged surface area (maximum of 196,350 m²) of Pool and Control for urban, natural, or agricultural land-use. Points are coloured by land-use category. Urban and agricultural land use together are defined as “land-use intensity”, which together with natural land-use is again the total area. Points are presented with a vertical jitter to allow overlapping values to be visible.*

*Table S1 Overview of study sites (Site), their geographic location (Latitude, Longitude), the year of sampling (Year), the approximated number of years since colonization of beavers (Age), the dominant habitat type (Habitat type), the stream section order classified as Mouth = 1, Middle = 2, and Upper = 3 (Stream section order), and the location of the respective control site, upstream or downstream (Control site)*

| **Site** | **Latitude** | **Longitude** | **Year** | **System Age** | **Habitat Type** | **Stream Section Order** | **Control Site** |
| --- | --- | --- | --- | --- | --- | --- | --- |
| Tegel | 47.55452 | 8.86189 | 2021 | 12 | open | 1 (Mouth) | downstream |
| Gile | 47.59317 | 8.75724 | 2021 | 5 | open | 2 (Middle) | upstream |
| Ellikon | 47.55402 | 8.80721 | 2021 | 8 | open | 2 (Middle) | upstream |
| Hemishofen | 47.68361 | 8.84713 | 2021 | 14 | open | 3 (Upper) | downstream |
| Nider | 47.60543 | 8.62667 | 2021 | 12 | Forest | 2 (Middle) | downstream |
| Logge | 47.62019 | 9.18029 | 2021 | 9 | Forest | 2 (Middle) | downstream |
| Rot | 47.08982 | 7.77282 | 2021 | 4 | Forest | 1 (Mouth) | downstream |
| Biber | 47.69095 | 8.81503 | 2021 | 5 | Forest | 2 (Middle) | upstream |
| Chriesbach | 47.408407 | 8.628318 | 2022 | 4 | open | 1 (Mouth) | downstream |
| Leugene | 47.169156 | 7.323065 | 2022 | 5 | open | 3 (Upper) | upstream |
| Gaebelbach | 46.946125 | 7.343173 | 2022 | 5 | open | 2 (Middle) | upstream |
| Weierbach | 47.165841 | 7.625691 | 2022 | 11 | open | 2 (Middle) | upstream |
| Haslibach | 46.994576 | 7.22699 | 2022 | 8 | Forest | 2 (Middle) | upstream |
| Riedgrabe | 47.204482 | 7.724176 | 2022 | 12 | Forest | 3 (Upper) | downstream |
| Coruz | 46.637545 | 6.683462 | 2022 | 11 | Forest | 3 (Upper) | downstream |
| Talent | 46.588417 | 6.681888 | 2022 | 5 | Forest | 3 (Upper) | upstream |

*Table S2: Overview of land-use types, their original German names, English translations, and land-use type assignments in the analyses. The land-use types were grouped into three main categories: agricultural, natural, and urban. Agricultural also includes biodiversity-friendly land-uses, such as extensively managed meadows, fallow land, and native hedgerows, for which farmers are entitled to compensation as well as crop fields, intensively managed pastures, etc. Natural areas encompass forests, wetlands, and water bodies, while urban areas include built-up land and recreational spaces.*

| Original German Name | English Translation | Land-use type |
| --- | --- | --- |
| **Extensiv genutzte Wiesen ohne Weiden** | Extensively used meadows without pastures | agriculture |
| **Bluehstreifen fuer Bestaeuber und andere Nuetzlinge** | Flower strips for pollinators and other beneficial organisms | agriculture |
| **Extensiv genutzte Weiden** | Extensively used pastures | agriculture |
| **Hochstamm Feldobstbaeume** | Standard field fruit trees | agriculture |
| **Uebrige unproduktive Flaechen z.B.gemulchte Flaechen stark verunkrautete Flaechen Hecken ohne Pufferstreifen** | Other unproductive areas (e.g, mulched areas, heavily overgrown areas, hedges) | agriculture |
| **Buntbrache** | Multicolored fallow land | agriculture |
| **Uebrige Gruenflaeche (Dauergruenflaeche), beitragsberechtigt** | Other green areas (permanent grassland), eligible for subsidies | agriculture |
| **Uebrige Dauerkulturen, beitragsberechtigt, aggregiert** | Other permanent crops, eligible for subsidies, aggregated | agriculture |
| **Kulturen in ganzjaehrig geschuetztem Anbau, beitragsberechtigt aggregiert** | Crops under year-round protected cultivation, eligible for subsidies, aggregated | agriculture |
| **Rotationsbrache** | Rotational fallow land | agriculture |
| **Wenig intensiv genutzte Wiesen (ohne Weiden)** | Low-intensity meadows (without pastures) | agriculture |
| **Streueflaechen in der LN** | Litter areas in agricultural zones | agriculture |
| **Hecken Feld und Ufergehölze mit Krautsaum** | Hedges, field, and riparian trees with herbaceous borders | agriculture |
| **Einheimische standortgerechte Einzelbaeume und Alleen** | Native site-appropriate solitary trees and avenues | agriculture |
| **Obstanlagen Aepfel** | Apple orchards | agriculture |
| **Obstanlagen Birnen** | Pear orchards | agriculture |
| **Obstanlagen Steinobst** | Stone fruit orchards | agriculture |
| **Silo und Gruenmais** | Silage and green maize | agriculture |
| **Zuckerrueben** | Sugar beets | agriculture |
| **Uebrige Dauerwiesen ohne Weiden** | Other permanent meadows without pastures | agriculture |
| **Kunstwiesen ohne Weiden** | Artificial meadows without pastures | agriculture |
| **Winterraps zur Speiseoelgewinnung** | Winter rapeseed for edible oil production | agriculture |
| **Winterweizen ohne Futterweizen der Sortenliste swiss granum** | Winter wheat (excluding feed wheat listed by Swiss granum) | agriculture |
| **Kartoffeln** | Potatoes | agriculture |
| **Koernermais** | Grain maize | agriculture |
| **Dinkel** | Spelt | agriculture |
| **Wintergerste** | Winter barley | agriculture |
| **Sommergerste** | Spring barley | agriculture |
| **Triticale** | Triticale | agriculture |
| **Sonnenblumen zur Speiseoelgewinnung** | Sunflowers for edible oil production | agriculture |
| **Gehoelzflaeche** | Wooded area | natural |
| **Feuchtgebiet** | Wetland | natural |
| **Fliessgewaesser** | Flowing water | natural |
| **Wald** | Forest | natural |
| **Wald nicht bestockt** | Unstocked forest | natural |
| **Stehende Gewaesser** | Standing water | natural |
| **Gebaeude** | Buildings | urban |
| **Abwasserreinigungsanlage** | Wastewater treatment plant | urban |
| **Sportplatzareal** | Sports facility area | urban |
| **Schul- und Hochschulareal** | School and university campus area | urban |
| **Schrebergartenareal** | Allotment garden area | urban |
| **Golfplatzareal** | Golf course area | urban |
| **Truppenuebungsplatz** | Military training area | urban |
| **Unbefestigte, natuerliche Wege** | Unpaved, natural paths | urban |
| **fehlend** | missing | urban |

*Table S3 Overview of the assignment of the operational taxonomic units (morphospecies) of plankton and assignment to taxonomic levels.*

| **Morphospecies** | **Kingdom** | **Phylum** | **Class** | **Order** | **Suborder** | **Family** | **Genus** |
| --- | --- | --- | --- | --- | --- | --- | --- |
| diatom | NA | Bacillariophyta | Bacillariophyceae | NA | NA | NA | NA |
| diatom_centric | NA | Bacillariophyta | Bacillariophyceae | NA | NA | NA | NA |
| diatom_chain | NA | Bacillariophyta | Bacillariophyceae | NA | NA | NA | NA |
| diatom_pennate_large | NA | Bacillariophyta | Bacillariophyceae | NA | NA | NA | NA |
| diatom_pennate_medium | NA | Bacillariophyta | Bacillariophyceae | NA | NA | NA | NA |
| diatom_pennate_small | NA | Bacillariophyta | Bacillariophyceae | NA | NA | NA | NA |
| pennate | NA | Bacillariophyta | NA | NA | NA | NA | NA |
| nitzschia | NA | Bacillariophyta | Bacillariophyceae | NA | NA | Bacillariaceae | Nitzschia |
| chaoborus | Metazoa | Arthropoda | Insecta | Diptera | Nematocera | Chaoboridae | Chaoborus |
| green_algae | Viridiplantae | NA | NA | NA | NA | NA | NA |
| green_algae_filament | Viridiplantae | NA | NA | NA | NA | NA | NA |
| chlorophyte | Viridiplantae | Chlorophyta | NA | NA | NA | NA | NA |
| chlorophyte_dividing | Viridiplantae | Chlorophyta | NA | NA | NA | NA | NA |
| chlorophyte_small | Viridiplantae | Chlorophyta | NA | NA | NA | NA | NA |
| ciliate | NA | Ciliophora | NA | NA | NA | NA | NA |
| ciliate_colony | NA | Ciliophora | NA | NA | NA | NA | NA |
| closterium | Viridiplantae | Streptophyta | Zygnemophyceae | Desmidiales | NA | Closteriaceae | Closterium |
| copepod | Metazoa | Arthropoda | Hexanauplia | NA | NA | NA | NA |
| nauplius | Metazoa | Arthropoda | NA | NA | NA | NA | NA |
| cryptomonas | NA | NA | Cryptophyceae | Cryptomonadales | NA | Cryptomonadaceae | Cryptomonas |
| cryptophyte | NA | NA | Cryptophyceae | NA | NA | NA | NA |
| cyanobacteria_filament | NA | Cyanobacteriota | NA | NA | NA | NA | NA |
| cyclops | Metazoa | Arthropoda | Hexanauplia | Cyclopoida | NA | Cyclopidae | Cyclops |
| daphnia | Metazoa | Arthropoda | Branchiopoda | Diplostraca | Cladocera | Daphniidae | Daphnia |
| dinobryon | NA | NA | Chrysophyceae | Chromulinales | NA | Dinobryaceae | Dinobryon |
| dinoflagelate | NA | NA | Dinophyceae | NA | NA | NA | NA |
| dinoflagellate_cyst | NA | NA | Dinophyceae | NA | NA | NA | NA |
| cladocera | Metazoa | Arthropoda | Branchiopoda | Diplostraca | NA | NA | NA |
| euglenoid | NA | Euglenozoa | Euglenida | NA | NA | NA | NA |
| gomphonema | NA | Bacillariophyta | Bacillariophyceae | Cymbellales | NA | Gomphonemataceae | Gomphonema |
| nematode | Metazoa | Nematoda | NA | NA | NA | NA | NA |
| phacus | NA | Euglenozoa | Euglenida | Euglenales | NA | Phacaceae | Phacus |
| rotifer | Metazoa | Rotifera | NA | NA | NA | NA | NA |
| scenedesmus | Viridiplantae | Chlorophyta | Chlorophyceae | Sphaeropleales | NA | Scenedesmaceae | Scenedesmus |
| staurastrum | Viridiplantae | Streptophyta | Zygnemophyceae | Desmidiales | NA | Desmidiaceae | Staurastrum |
| synura | NA | NA | Synurophyceae | Synurales | NA | Mallomonadaceae | Synura |
| uroglena | NA | NA | Chrysophyceae | Chromulinales | NA | Chromulinaceae | Uroglena |
| jochalgen | Viridiplantae | Streptophyta | Zygnemophyceae | Zygnematales | NA | NA | NA |
| larvae | Metazoa | Arthropoda | Insecta | NA | NA | NA | NA |
| heliozoa | NA | NA | NA | NA | NA | NA | NA |
| phytoplankton | NA | NA | NA | NA | NA | NA | NA |
| phytoplankton_colony | NA | NA | NA | NA | NA | NA | NA |
| phytoplankton_filament | NA | NA | NA | NA | NA | NA | NA |
| plankton | NA | NA | NA | NA | NA | NA | NA |
| protist | NA | NA | NA | NA | NA | NA | NA |
| worm | NA | NA | NA | NA | NA | NA | NA |
| zooplankton | NA | NA | NA | NA | NA | NA | NA |
| zooplankton_filament | NA | NA | NA | NA | NA | NA | NA |

*Table S4: GLMM results for assessing the differences in species richness, abundance, unique species richness and evenness between Pool and Control areas for each aquatic and terrestrial taxa. estimate = estimated value of Intercept (equivalent to the Control) and Pool, std. error = Standard error, p-value = indicating significant differences.*

| **Metric** | **Community** | **Estimate** | **Std.error** | **p.value** |
| --- | --- | --- | --- | --- |
| Richness | amphibians | 2.05 | 0.53 | **<0.001** |
| Richness | aquatic invertebrates | -0.02 | 0.09 | 0.82 |
| Richness | flying arthropods | -0.04 | 0.14 | 0.78 |
| Richness | terrestrial invertebrates | -0.08 | 0.1 | 0.44 |
| Richness | bats | 0.2 | 0.15 | 0.18 |
| Richness | dragonflies | 0.81 | 0.15 | **<0.001** |
| Richness | fishes | 0.44 | 0.22 | **<0.05** |
| Richness | macrophytes | 1.06 | 0.19 | **<0.001** |
| Richness | plankton | 0.36 | 0.15 | **<0.05** |
| Richness | terrestrial plants | 0.28 | 0.11 | **<0.05** |
| Abundance | amphibians | 6.49 | 0.32 | **<0.001** |
| Abundance | aquatic invertebrates | 0 | 0.23 | 0.99 |
| Abundance | flying arthropods | 0.15 | 0.08 | 0.08 |
| Abundance | terrestrial invertebrates | 0.48 | 0.31 | 0.12 |
| Abundance | bats | 0.45 | 0.27 | 0.1 |
| Abundance | dragonflies | 1.2 | 0.41 | **<0.01** |
| Abundance | fishes | 0.78 | 0.46 | 0.09 |
| Abundance | macrophytes | 0.8 | 0.39 | **<0.05** |
| Abundance | plankton | 0.74 | 0.29 | **<0.05** |
| Abundance | terrestrial plants | -0.15 | 0.14 | 0.29 |
| Unique Species | amphibians | 22.97 | 3.87 | **<0.001** |
| Unique Species | aquatic invertebrates | -0.04 | 0.17 | 0.82 |
| Unique Species | flying arthropods | -0.27 | 0.37 | 0.46 |
| Unique Species | terrestrial invertebrates | -0.37 | 0.26 | 0.16 |
| Unique Species | bats | 1.39 | 0.47 | **<0.01** |
| Unique Species | dragonflies | 2.49 | 0.39 | **<0.001** |
| Unique Species | fishes | 1.57 | 0.49 | **<0.01** |
| Unique Species | macrophytes | 2.73 | 0.37 | **<0.001** |
| Unique Species | plankton | 0.82 | 0.29 | **<0.01** |
| Unique Species | terrestrial plants | 0.36 | 0.12 | **<0.01** |
| Evenness | amphibians | 0.48 | 0.4 | 0.23 |
| Evenness | aquatic invertebrates | 0.05 | 0.22 | 0.83 |
| Evenness | flying arthropods | -0.21 | 0.09 | **<0.05** |
| Evenness | terrestrial invertebrates | -0.3 | 0.25 | 0.23 |
| Evenness | bats | -0.21 | 0.17 | 0.22 |
| Evenness | dragonflies | 0.64 | 0.39 | 0.1 |
| Evenness | fishes | 0.15 | 0.32 | 0.65 |
| Evenness | macrophytes | 0.69 | 0.42 | 0.09 |
| Evenness | plankton | 0.3 | 0.43 | 0.5 |
| Evenness | terrestrial plants | -0.02 | 0.21 | 0.91 |

*Table S5 Dharma results: Model selection was guided by AIC and DHARMa diagnostics, with R2 to support. Poisson GLMMs were generally sufficient for species richness and unique species richness, although negative binomial (nb) models improved fit for some taxa (macrophytes for richness, aquatic invertebrates for unique species richness). Abundance models typically required negative binomial distributions, except for amphibians (Poisson) and macrophytes, where a zero-inflated negative binomial model was used due to strong overdispersion and excess zeros. Although dispersion tests indicated underdispersion in some species richness poisson models (terrestrial invertebrates, flying arthropods, bats), changing to negative binomial models did not improve model fit based on AIC or residual diagnostics. Thus, Poisson GLMMs were retained in these cases for parsimony and interpretability. However, these results need to be interpreted with caution. All other final models passed DHARMa diagnostic checks (p > 0.05) for dispersion, zero-inflation, outliers, and residual uniformity. Highlighted in* ***bold*** *are where models deviate from expectations. Part A has an overview of model diagnostics, including the metric analyzed (Metric), the biological community (Community), the initial assumed model distribution (Base Model), results from dispersion tests (Dispersion, Dispersion_p), zero-inflation (Zero_inflation_p), outliers (Outlier_p) and residual uniformity (Uniformity_p) as well as the final model choice (Final Decision), and relevant notes if the final model deviated from the base model (Comments). Part B has an overview of model comparison results with the metric analyzed (Metric), the biological community (Community), the initial model distribution (Base Model), AIC values for Poisson and negative binomial models (AIC_poisson, AIC_neg binomial), their difference (Delta_AIC), marginal and conditional R² values for both models (R2m_poisson, R2m_nb, R2c_poisson, R2c_nb), R² differences (Delta R2m, Delta R2c), AIC-based model preference (Decision_AIC), and best model based on R² (Best Model_R2).*

| A) |  |  |  |  |  |  |  |  |  |
| --- | --- | --- | --- | --- | --- | --- | --- | --- | --- |
| Metric | **Community** | **Base Model** | **Dispersion** | **Dispersion_p** | **Zero_ inflation_p** | **Outlier_p** | **Uniformity_p** | **Final Decision** | **Comments** |
| richness | amphibians | poisson | 0.86 | 0.98 | 1.00 | 1.00 | 0.90 | poisson |  |
| richness | aquatic_invertebrates | poisson | 1.03 | 0.84 | 1.00 | 1.00 | 0.80 | poisson | R2 meaningless difference |
| richness | arthropods_terrestrial | poisson | **0.52** | **0.02** | 1.00 | 1.00 | 0.65 | poisson |  |
| richness | dragonflies | poisson | 0.78 | 0.91 | 0.73 | 1.00 | 0.94 | poisson |  |
| richness | fish | poisson | 0.98 | 0.70 | 1.00 | 1.00 | 0.80 | poisson | R2 meaningless difference |
| richness | macrophytes | poisson | 1.26 | 0.38 | **0.02** | 1.00 | 0.52 | **nb** | nb overall better, AIC and solves zero-inflation of model |
| richness | plankton | poisson | 0.88 | 0.88 | 0.93 | 1.00 | 0.87 | poisson |  |
| richness | plants_terrestrial | poisson | 0.89 | 0.94 | 1.00 | 1.00 | 0.94 | poisson |  |
| richness | arthropods_flying | poisson | **0.26** | **0.00** | 1.00 | 1.00 | 0.37 | poisson | R2 meaningless difference |
| richness | bats | poisson | **0.34** | **0.02** | 1.00 | 1.00 | 0.32 | poisson |  |
| abundance | amphibians | nb | 0.00 | 0.31 | 0.59 | 1.00 | 0.37 | **poisson** | no significant gain for nb, which even has a failed levene test for homogeneity of variance in dharma, so we take poisson. R2 meaningless difference |
| abundance | aquatic_invertebrates | nb | 0.94 | 0.94 | 1.00 | 1.00 | 0.91 | nb | R2 meaningless difference |
| abundance | arthropods_terrestrial | nb | 1.01 | 0.53 | 1.00 | 1.00 | 0.91 | nb | AIC difference overules R2 |
| abundance | dragonflies | nb | 0.75 | 0.88 | 0.42 | 1.00 | 0.96 | nb | AIC difference overules R2 |
| abundance | fish | nb | 0.03 | 0.98 | 0.10 | 1.00 | 0.23 | nb | AIC difference overules R2 |
| abundance | macrophytes | nb | 0.46 | 0.88 | **0.00** | **0.02** | 0.17 | **zero-inflated** | zero-inflated in the end (not in this table) |
| abundance | plankton | nb | 0.29 | 0.94 | 0.34 | 1.00 | 0.75 | nb | AIC difference overules R2 |
| abundance | plants_terrestrial | nb | 0.70 | 0.36 | 1.00 | 0.16 | 0.21 | nb | AIC difference overules R2 |
| abundance | arthropods_flying | nb | 0.75 | 0.95 | 1.00 | 1.00 | 0.54 | nb | AIC difference overules R2 |
| abundance | bats | nb | 1.11 | 0.64 | 1.00 | 1.00 | 0.99 | nb | AIC difference overules R2 |
| unique_species | amphibians | poisson | 1.06 | 0.79 | 1.00 | 1.00 | 0.32 | poisson | R2 meaningless difference |
| unique_species | aquatic_invertebrates | poisson | 1.56 | 0.10 | 1.00 | 1.00 | 0.52 | **nb** | follow AIC, also model diagnostics better |
| unique_species | arthropods_terrestrial | poisson | 1.44 | 0.14 | 0.54 | 1.00 | 0.91 | poisson |  |
| unique_species | dragonflies | poisson | 1.10 | 0.64 | 0.83 | 1.00 | 0.80 | poisson | R2 meaningless difference |
| unique_species | fish | poisson | 0.90 | 0.99 | 0.90 | 1.00 | 0.80 | poisson |  |
| unique_species | macrophytes | poisson | 1.29 | 0.46 | 0.94 | 1.00 | 0.25 | poisson | R2 meaningless difference |
| unique_species | plankton | poisson | 1.27 | 0.43 | 0.38 | 1.00 | 0.77 | poisson |  |
| unique_species | plants_terrestrial | poisson | 0.81 | 0.57 | 1.00 | 1.00 | 0.67 | poisson |  |
| unique_species | arthropods_flying | poisson | 1.08 | 0.79 | 1.00 | 1.00 | 0.58 | poisson |  |
| unique_species | bats | poisson | 1.25 | 0.47 | 1.00 | 1.00 | 0.90 | poisson |  |

| B) |  |  |  |  |  |  |  |  |  |  |  |  |  |
| --- | --- | --- | --- | --- | --- | --- | --- | --- | --- | --- | --- | --- | --- |
| Metric | **Community** | **Base Model** | **AIC_ poisson** | **AIC_neg binomial** | **Delta_AIC** | **R2m_ poisson** | **R2m_nb** | **R2c_ poisson** | **R2c_nb** | **Delta R2m** | **Delta R2c** | **Decision_AIC** | **Best Model_R2** |
| richness | amphibians | poisson | 81.17 | 83.17 | -2.00 | 0.49 | 0.49 | 0.56 | 0.56 | 0.00 | 0.00 | poisson | poisson |
| richness | aquatic_invertebrates | poisson | 199.65 | 201.65 | -2.00 | 0.00 | 0.00 | 0.00 | 0.00 | 0.00 | 0.00 | poisson | **nb** |
| richness | arthropods_terrestrial | poisson | 159.04 | 161.04 | -2.00 | 0.02 | 0.02 | 0.02 | 0.02 | 0.00 | 0.00 | poisson | poisson |
| richness | dragonflies | poisson | 171.37 | 173.37 | -2.00 | 0.26 | 0.26 | 0.27 | 0.27 | 0.00 | 0.00 | poisson | poisson |
| richness | fish | poisson | 129.61 | 131.61 | -2.00 | 0.06 | 0.06 | 0.06 | 0.06 | 0.00 | 0.00 | poisson | **nb** |
| richness | macrophytes | poisson | 181.58 | 179.85 | **1.73** | 0.63 | 0.53 | 0.64 | 0.56 | 0.10 | 0.08 | **nb** | **poisson** |
| richness | plankton | poisson | 162.32 | 164.32 | -2.00 | 0.08 | 0.08 | 0.08 | 0.08 | 0.00 | 0.00 | poisson | poisson |
| richness | plants_terrestrial | poisson | 180.75 | 182.75 | -2.00 | 0.09 | 0.09 | 0.09 | 0.09 | 0.00 | 0.00 | poisson | poisson |
| richness | arthropods_flying | poisson | 79.38 | 81.38 | -2.00 | 0.01 | 0.01 | 0.01 | 0.01 | 0.00 | 0.00 | poisson | **nb** |
| richness | bats | poisson | 78.85 | 80.85 | -2.00 | 0.11 | 0.11 | 0.11 | 0.11 | 0.00 | 0.00 | poisson | poisson |
| abundance | amphibians | nb | 233.49 | 235.47 | **1.98** | 0.47 | 0.47 | 0.47 | 0.47 | 0.00 | 0.00 | **poisson** | **poisson** |
| abundance | aquatic_invertebrates | nb | 2152.57 | 412.16 | -1740.41 | 0.00 | 0.00 | 0.00 | 0.00 | 0.00 | 0.00 | nb | **poisson** |
| abundance | arthropods_terrestrial | nb | 5241.27 | 457.16 | -4784.11 | 0.09 | 0.06 | 0.09 | 0.07 | -0.04 | -0.03 | nb | **poisson** |
| abundance | dragonflies | nb | 1478.47 | 364.27 | -1114.20 | 0.32 | 0.22 | 0.32 | 0.31 | -0.10 | -0.01 | nb | **poisson** |
| abundance | fish | nb | 4513.20 | 395.34 | -4117.86 | 0.06 | 0.03 | 0.06 | 0.03 | -0.03 | -0.03 | nb | **poisson** |
| abundance | macrophytes | nb | 9309.18 | 9148.59 | -160.59 | 0.18 | 0.18 | 0.18 | 0.18 | 0.00 | 0.00 | nb | nb |
| abundance | plankton | nb | 574.82 | 282.50 | -292.32 | 0.08 | 0.06 | 0.08 | 0.06 | -0.02 | -0.02 | nb | **poisson** |
| abundance | plants_terrestrial | nb | 457.77 | 335.38 | -122.39 | 0.06 | 0.04 | 0.06 | 0.04 | -0.02 | -0.02 | nb | **poisson** |
| abundance | arthropods_flying | nb | 259.26 | 203.65 | -55.61 | 0.04 | 0.03 | 0.04 | 0.03 | 0.00 | 0.00 | nb | **poisson** |
| abundance | bats | nb | 3160.29 | 267.25 | -2893.04 | 0.19 | 0.13 | 0.19 | 0.14 | -0.06 | -0.05 | nb | **poisson** |
| unique_species | amphibians | poisson | 58.11 | 60.11 | -2.00 | 0.99 | 0.99 | 0.99 | 0.99 | 0.00 | 0.00 | poisson | **nb** |
| unique_species | aquatic_invertebrates | poisson | 192.85 | 187.86 | **4.99** | 0.00 | 0.00 | 0.01 | 0.00 | 0.00 | 0.00 | **nb** | poisson |
| unique_species | arthropods_terrestrial | poisson | 132.26 | 132.70 | -0.44 | 0.08 | 0.06 | 0.10 | 0.08 | 0.02 | 0.02 | poisson | poisson |
| unique_species | dragonflies | poisson | 119.67 | 121.55 | -1.88 | 0.75 | 0.75 | 0.76 | 0.77 | 0.00 | 0.00 | poisson | **nb** |
| unique_species | fish | poisson | 80.21 | 82.21 | -2.00 | 0.24 | 0.24 | 0.30 | 0.30 | 0.00 | 0.00 | poisson | poisson |
| unique_species | macrophytes | poisson | 132.30 | 133.98 | -1.68 | 0.82 | 0.82 | 0.83 | 0.83 | 0.00 | 0.00 | poisson | **nb** |
| unique_species | plankton | poisson | 118.04 | 119.91 | -1.87 | 0.25 | 0.25 | 0.28 | 0.28 | 0.00 | 0.00 | poisson | poisson |
| unique_species | plants_terrestrial | poisson | 163.75 | 165.75 | -2.00 | 0.19 | 0.19 | 0.20 | 0.20 | 0.00 | 0.00 | poisson | poisson |
| unique_species | arthropods_flying | poisson | 58.59 | 60.59 | -2.00 | 0.04 | 0.04 | 0.04 | 0.04 | 0.00 | 0.00 | poisson | poisson |
| unique_species | bats | poisson | 55.61 | 57.54 | -1.93 | 0.49 | 0.47 | 0.55 | 0.53 | 0.02 | 0.02 | poisson | poisson |

*Table S6 Linear model results for testing the effect of urban, agricultural and natural land-use on changes in biodiversity metrics species richness, abundance and evenness (Pool − Control) for the different aquatic and terrestrial taxa. Estimate = estimated effect size of land-use on* Δ *richness, Std. Error = standard error of the estimate, R² and adj. R² = model fit indicators, and p-value = significance of the relationship. Human is the combined effect of urban and agricultural cover, called land-use intensity in the main text.* Δ *unique species richness is the same as richness and therefore model results are not shown separately.*

| **Metric** | **Community** | **Land Use** | **Estimate** | **Std.error** | **R.squared** | **Adj.r.squared** | **p.value** |
| --- | --- | --- | --- | --- | --- | --- | --- |
| richness | amphibians | agriculture | -1.74E-05 | 5.64E-06 | 0.41 | 0.36 | 0.008 |
| richness | aquatic invertebrates | agriculture | -2.07E-05 | 2.91E-05 | 0.03 | -0.03 | 0.490 |
| richness | flying arthropods | agriculture | 3.26E-05 | 1.50E-05 | 0.44 | 0.35 | 0.073 |
| richness | terrestrial invertebrates | agriculture | 3.29E-06 | 1.84E-05 | 0.00 | -0.07 | 0.861 |
| richness | bats | agriculture | -1.08E-05 | 2.10E-05 | 0.04 | -0.12 | 0.625 |
| richness | dragonflies | agriculture | -4.54E-05 | 1.56E-05 | 0.38 | 0.33 | 0.012 |
| richness | fish | agriculture | 9.36E-07 | 6.66E-06 | 0.00 | -0.07 | 0.890 |
| richness | macrophytes | agriculture | -4.84E-05 | 2.43E-05 | 0.22 | 0.17 | 0.066 |
| richness | plankton | agriculture | -2.12E-05 | 1.56E-05 | 0.12 | 0.05 | 0.195 |
| richness | terrestrial plants | agriculture | 2.49E-06 | 1.54E-05 | 0.00 | -0.07 | 0.874 |
| richness | amphibians | urban | -2.23E-05 | 1.20E-05 | 0.20 | 0.14 | 0.085 |
| richness | aquatic invertebrates | urban | 1.28E-05 | 5.43E-05 | 0.00 | -0.07 | 0.817 |
| richness | flying arthropods | urban | 2.42E-05 | 2.40E-05 | 0.15 | 0.00 | 0.352 |
| richness | terrestrial invertebrates | urban | 7.00E-05 | 2.82E-05 | 0.31 | 0.26 | 0.026 |
| richness | bats | urban | -3.00E-05 | 2.49E-05 | 0.19 | 0.06 | 0.274 |
| richness | dragonflies | urban | -4.54E-05 | 3.42E-05 | 0.11 | 0.05 | 0.206 |
| richness | fish | urban | 1.60E-06 | 1.22E-05 | 0.00 | -0.07 | 0.898 |
| richness | macrophytes | urban | -8.47E-05 | 4.51E-05 | 0.20 | 0.14 | 0.081 |
| richness | plankton | urban | -5.39E-06 | 3.03E-05 | 0.00 | -0.07 | 0.862 |
| richness | terrestrial plants | urban | 4.42E-05 | 2.57E-05 | 0.17 | 0.12 | 0.107 |
| richness | amphibians | natural | 1.44E-05 | 4.15E-06 | 0.46 | 0.42 | 0.004 |
| richness | aquatic invertebrates | natural | 1.01E-05 | 2.28E-05 | 0.01 | -0.06 | 0.664 |
| richness | flying arthropods | natural | -2.41E-05 | 1.04E-05 | 0.47 | 0.38 | 0.061 |
| richness | terrestrial invertebrates | natural | -1.44E-05 | 1.37E-05 | 0.07 | 0.01 | 0.311 |
| richness | bats | natural | 1.47E-05 | 1.41E-05 | 0.15 | 0.01 | 0.338 |
| richness | dragonflies | natural | 3.53E-05 | 1.21E-05 | 0.38 | 0.33 | 0.011 |
| richness | fish | natural | -8.45E-07 | 5.15E-06 | 0.00 | -0.07 | 0.872 |
| richness | macrophytes | natural | 4.41E-05 | 1.77E-05 | 0.31 | 0.26 | 0.026 |
| richness | plankton | natural | 1.37E-05 | 1.23E-05 | 0.08 | 0.02 | 0.285 |
| richness | terrestrial plants | natural | -9.37E-06 | 1.17E-05 | 0.04 | -0.02 | 0.436 |
| richness | amphibians | human | -1.44E-05 | 4.15E-06 | 0.46 | 0.42 | 0.004 |
| richness | aquatic invertebrates | human | -1.01E-05 | 2.28E-05 | 0.01 | -0.06 | 0.664 |
| richness | flying arthropods | human | 2.41E-05 | 1.04E-05 | 0.47 | 0.38 | 0.061 |
| richness | terrestrial invertebrates | human | 1.44E-05 | 1.37E-05 | 0.07 | 0.01 | 0.311 |
| richness | bats | human | -1.47E-05 | 1.41E-05 | 0.15 | 0.01 | 0.338 |
| richness | dragonflies | human | -3.53E-05 | 1.21E-05 | 0.38 | 0.33 | 0.011 |
| richness | fish | human | 8.45E-07 | 5.15E-06 | 0.00 | -0.07 | 0.872 |
| richness | macrophytes | human | -4.41E-05 | 1.77E-05 | 0.31 | 0.26 | 0.026 |
| richness | plankton | human | -1.37E-05 | 1.23E-05 | 0.08 | 0.02 | 0.285 |
| richness | terrestrial plants | human | 9.37E-06 | 1.17E-05 | 0.04 | -0.02 | 0.436 |
| abundance | amphibians | agriculture | -7.04E-03 | 2.13E-03 | 0.44 | 0.40 | 0.005 |
| abundance | aquatic invertebrates | agriculture | -7.01E-04 | 1.31E-03 | 0.02 | -0.05 | 0.600 |
| abundance | flying arthropods | agriculture | 1.44E-04 | 8.45E-04 | 0.00 | -0.16 | 0.870 |
| abundance | terrestrial invertebrates | agriculture | -4.39E-03 | 3.63E-03 | 0.09 | 0.03 | 0.247 |
| abundance | bats | agriculture | -7.36E-03 | 1.04E-02 | 0.08 | -0.08 | 0.506 |
| abundance | dragonflies | agriculture | -8.34E-04 | 9.63E-04 | 0.05 | -0.02 | 0.401 |
| abundance | fish | agriculture | 3.84E-03 | 5.41E-03 | 0.03 | -0.03 | 0.490 |
| abundance | macrophytes | agriculture | -1.00E-02 | 4.69E-03 | 0.25 | 0.19 | 0.050 |
| abundance | plankton | agriculture | 9.88E-05 | 3.58E-04 | 0.01 | -0.07 | 0.787 |
| abundance | terrestrial plants | agriculture | 8.44E-05 | 2.37E-04 | 0.01 | -0.06 | 0.727 |
| abundance | amphibians | urban | -1.11E-02 | 4.29E-03 | 0.32 | 0.27 | 0.022 |
| abundance | aquatic invertebrates | urban | -1.83E-03 | 2.37E-03 | 0.04 | -0.03 | 0.453 |
| abundance | flying arthropods | urban | 3.01E-04 | 1.09E-03 | 0.01 | -0.15 | 0.792 |
| abundance | terrestrial invertebrates | urban | -5.30E-04 | 7.00E-03 | 0.00 | -0.07 | 0.941 |
| abundance | bats | urban | -9.34E-03 | 1.35E-02 | 0.07 | -0.08 | 0.515 |
| abundance | dragonflies | urban | -2.82E-03 | 1.65E-03 | 0.17 | 0.11 | 0.110 |
| abundance | fish | urban | -2.72E-04 | 1.01E-02 | 0.00 | -0.07 | 0.979 |
| abundance | macrophytes | urban | -1.28E-02 | 9.31E-03 | 0.12 | 0.06 | 0.190 |
| abundance | plankton | urban | 7.30E-04 | 6.30E-04 | 0.09 | 0.02 | 0.266 |
| abundance | terrestrial plants | urban | 3.03E-04 | 4.30E-04 | 0.03 | -0.03 | 0.492 |
| abundance | amphibians | natural | 6.19E-03 | 1.45E-03 | 0.57 | 0.54 | 0.001 |
| abundance | aquatic invertebrates | natural | 7.46E-04 | 1.00E-03 | 0.04 | -0.03 | 0.469 |
| abundance | flying arthropods | natural | -1.66E-04 | 6.02E-04 | 0.01 | -0.15 | 0.793 |
| abundance | terrestrial invertebrates | natural | 2.73E-03 | 2.86E-03 | 0.06 | -0.01 | 0.357 |
| abundance | bats | natural | 6.62E-03 | 7.26E-03 | 0.12 | -0.02 | 0.397 |
| abundance | dragonflies | natural | 1.00E-03 | 7.17E-04 | 0.12 | 0.06 | 0.184 |
| abundance | fish | natural | -2.25E-03 | 4.22E-03 | 0.02 | -0.05 | 0.602 |
| abundance | macrophytes | natural | 8.30E-03 | 3.55E-03 | 0.28 | 0.23 | 0.035 |
| abundance | plankton | natural | -1.89E-04 | 2.73E-04 | 0.03 | -0.04 | 0.501 |
| abundance | terrestrial plants | natural | -1.05E-04 | 1.82E-04 | 0.02 | -0.05 | 0.576 |
| abundance | amphibians | human | -6.19E-03 | 1.45E-03 | 0.57 | 0.54 | 0.001 |
| abundance | aquatic invertebrates | human | -7.46E-04 | 1.00E-03 | 0.04 | -0.03 | 0.469 |
| abundance | flying arthropods | human | 1.66E-04 | 6.02E-04 | 0.01 | -0.15 | 0.793 |
| abundance | terrestrial invertebrates | human | -2.73E-03 | 2.86E-03 | 0.06 | -0.01 | 0.357 |
| abundance | bats | human | -6.62E-03 | 7.26E-03 | 0.12 | -0.02 | 0.397 |
| abundance | dragonflies | human | -1.00E-03 | 7.17E-04 | 0.12 | 0.06 | 0.184 |
| abundance | fish | human | 2.25E-03 | 4.22E-03 | 0.02 | -0.05 | 0.602 |
| abundance | macrophytes | human | -8.30E-03 | 3.55E-03 | 0.28 | 0.23 | 0.035 |
| abundance | plankton | human | 1.89E-04 | 2.73E-04 | 0.03 | -0.04 | 0.501 |
| abundance | terrestrial plants | human | 1.05E-04 | 1.82E-04 | 0.02 | -0.05 | 0.576 |
| evenness | amphibians | agriculture | -2.66E-06 | 1.92E-06 | 0.12 | 0.06 | 0.188 |
| evenness | aquatic invertebrates | agriculture | -2.34E-07 | 1.12E-06 | 0.00 | -0.07 | 0.837 |
| evenness | flying arthropods | agriculture | 8.00E-07 | 4.80E-07 | 0.32 | 0.20 | 0.147 |
| evenness | terrestrial invertebrates | agriculture | 1.57E-06 | 1.41E-06 | 0.08 | 0.02 | 0.286 |
| evenness | bats | agriculture | -5.99E-07 | 1.13E-06 | 0.04 | -0.11 | 0.615 |
| evenness | dragonflies | agriculture | 5.11E-07 | 2.25E-06 | 0.00 | -0.07 | 0.823 |
| evenness | fish | agriculture | -1.58E-07 | 1.20E-06 | 0.00 | -0.07 | 0.897 |
| evenness | macrophytes | agriculture | 1.55E-07 | 2.33E-06 | 0.00 | -0.07 | 0.948 |
| evenness | plankton | agriculture | 1.21E-06 | 2.36E-06 | 0.02 | -0.05 | 0.616 |
| evenness | terrestrial plants | agriculture | 1.07E-06 | 9.85E-07 | 0.08 | 0.01 | 0.298 |
| evenness | amphibians | urban | 3.22E-06 | 3.66E-06 | 0.05 | -0.02 | 0.393 |
| evenness | aquatic invertebrates | urban | 6.06E-07 | 2.04E-06 | 0.01 | -0.06 | 0.771 |
| evenness | flying arthropods | urban | -5.70E-07 | 7.15E-07 | 0.10 | -0.05 | 0.456 |
| evenness | terrestrial invertebrates | urban | 5.47E-07 | 2.70E-06 | 0.00 | -0.07 | 0.842 |
| evenness | bats | urban | 1.23E-06 | 1.41E-06 | 0.11 | -0.04 | 0.417 |
| evenness | dragonflies | urban | 1.48E-06 | 4.11E-06 | 0.01 | -0.06 | 0.724 |
| evenness | fish | urban | -1.52E-06 | 2.16E-06 | 0.03 | -0.04 | 0.494 |
| evenness | macrophytes | urban | -6.40E-06 | 3.91E-06 | 0.16 | 0.10 | 0.124 |
| evenness | plankton | urban | -2.49E-06 | 4.32E-06 | 0.02 | -0.05 | 0.574 |
| evenness | terrestrial plants | urban | -7.01E-07 | 1.87E-06 | 0.01 | -0.06 | 0.714 |
| evenness | amphibians | natural | 1.02E-06 | 1.56E-06 | 0.03 | -0.04 | 0.524 |
| evenness | aquatic invertebrates | natural | 3.21E-08 | 8.65E-07 | 0.00 | -0.07 | 0.971 |
| evenness | flying arthropods | natural | -2.35E-07 | 4.04E-07 | 0.05 | -0.10 | 0.582 |
| evenness | terrestrial invertebrates | natural | -1.04E-06 | 1.11E-06 | 0.06 | -0.01 | 0.365 |
| evenness | bats | natural | -6.92E-08 | 8.27E-07 | 0.00 | -0.17 | 0.936 |
| evenness | dragonflies | natural | -5.70E-07 | 1.74E-06 | 0.01 | -0.06 | 0.748 |
| evenness | fish | natural | 3.65E-07 | 9.23E-07 | 0.01 | -0.06 | 0.699 |
| evenness | macrophytes | natural | 1.05E-06 | 1.78E-06 | 0.02 | -0.05 | 0.566 |
| evenness | plankton | natural | -2.83E-07 | 1.84E-06 | 0.00 | -0.07 | 0.880 |
| evenness | terrestrial plants | natural | -5.14E-07 | 7.82E-07 | 0.03 | -0.04 | 0.522 |
| evenness | amphibians | human | -1.02E-06 | 1.56E-06 | 0.03 | -0.04 | 0.524 |
| evenness | aquatic invertebrates | human | -3.21E-08 | 8.65E-07 | 0.00 | -0.07 | 0.971 |
| evenness | flying arthropods | human | 2.35E-07 | 4.04E-07 | 0.05 | -0.10 | 0.582 |
| evenness | terrestrial invertebrates | human | 1.04E-06 | 1.11E-06 | 0.06 | -0.01 | 0.365 |
| evenness | bats | human | 6.92E-08 | 8.27E-07 | 0.00 | -0.17 | 0.936 |
| evenness | dragonflies | human | 5.70E-07 | 1.74E-06 | 0.01 | -0.06 | 0.748 |
| evenness | fish | human | -3.65E-07 | 9.23E-07 | 0.01 | -0.06 | 0.699 |
| evenness | macrophytes | human | -1.05E-06 | 1.78E-06 | 0.02 | -0.05 | 0.566 |
| evenness | plankton | human | 2.83E-07 | 1.84E-06 | 0.00 | -0.07 | 0.880 |
| evenness | terrestrial plants | human | 5.14E-07 | 7.82E-07 | 0.03 | -0.04 | 0.522 |

*Table S7 Overview of cumulative unique species for Pool and Control areas and to which taxa the respective species belongs*

| **Cumulative Unique Species in Pool** | **community** | **Cumulative Unique Species in Control** | **community** |
| --- | --- | --- | --- |
| Ichthyosaura alpestris | amphibians | Agabus | aquatic_invertebrates |
| Lissotriton helveticus | amphibians | Alainites muticus | aquatic_invertebrates |
| Aeshna cyanea | aquatic_invertebrates | Anacaena | aquatic_invertebrates |
| Agapetus | aquatic_invertebrates | Baetis melanonyx | aquatic_invertebrates |
| Allogamus auricollis | aquatic_invertebrates | Centroptilum luteolum | aquatic_invertebrates |
| Anabolia nervosa | aquatic_invertebrates | Cordulegaster bidentata | aquatic_invertebrates |
| Athripsodes albifrons | aquatic_invertebrates | Enochrus testaceus | aquatic_invertebrates |
| Athripsodes bilineatus | aquatic_invertebrates | Epeorus assimilis | aquatic_invertebrates |
| Baetis buceratus | aquatic_invertebrates | Galba truncatula | aquatic_invertebrates |
| Baetis vardarensis | aquatic_invertebrates | Gyraulus albus | aquatic_invertebrates |
| Brachycentrus subnubilus | aquatic_invertebrates | Habroleptoides confusa | aquatic_invertebrates |
| Caenis beskidensis | aquatic_invertebrates | Habrophlebia fusca | aquatic_invertebrates |
| Coenagrion puella | aquatic_invertebrates | Helochaeres lividus | aquatic_invertebrates |
| Corixa punctata | aquatic_invertebrates | Limnius germanus | aquatic_invertebrates |
| Ecnomus | aquatic_invertebrates | Limnius perrisi | aquatic_invertebrates |
| Elmis rietscheli | aquatic_invertebrates | Nemoura | aquatic_invertebrates |
| Haliplus lineatocollis | aquatic_invertebrates | Nemurella pictetii | aquatic_invertebrates |
| Hydropsyche angustipennis | aquatic_invertebrates | Orthetrum albistylum | aquatic_invertebrates |
| Hyphydrus ovatus | aquatic_invertebrates | Planorbarius corneus | aquatic_invertebrates |
| Leuctra geniculata | aquatic_invertebrates | Platycnemis pennipes | aquatic_invertebrates |
| Limnephilus germanus | aquatic_invertebrates | Plectrocnemia conspersa | aquatic_invertebrates |
| Limnephilus stigma | aquatic_invertebrates | Plectrocnemia geniculata | aquatic_invertebrates |
| Lymnaea stagnalis | aquatic_invertebrates | Polycentropus flavomaculatus | aquatic_invertebrates |
| Melampophylax mucoreus | aquatic_invertebrates | Potamophylax cingulatus | aquatic_invertebrates |
| Microvelia | aquatic_invertebrates | Procloeon bifidum | aquatic_invertebrates |
| Nepa cinerea | aquatic_invertebrates | Rhithrogena semicolorata | aquatic_invertebrates |
| Notonecta glauca | aquatic_invertebrates | Rhyacophila tristis | aquatic_invertebrates |
| Onychogomphus forcipatus | aquatic_invertebrates | Riolus cupreus | aquatic_invertebrates |
| Physidae | aquatic_invertebrates | Sericostoma | aquatic_invertebrates |
| Proasellus meridianus | aquatic_invertebrates | Orthoptera | arthropods_flying |
| Protonemura | aquatic_invertebrates | Plecoptera | arthropods_flying |
| Rhithrogena beskidensis | aquatic_invertebrates | Dermaptera | arthropods_terrestrial |
| Segmentina nitida | aquatic_invertebrates | Enchytraeida | arthropods_terrestrial |
| Sialis lutaria | aquatic_invertebrates | Orthoptera | arthropods_terrestrial |
| Somatochlora metallica | aquatic_invertebrates | Perca_fluviatilis | fish |
| Stenelmis | aquatic_invertebrates | chlorophyte | plankton |
| Ixodida | arthropods_terrestrial | chlorophyte_dividing | plankton |
| Zygentoma | arthropods_terrestrial | cyanobacteria_filament | plankton |
| Plecotus auritus | bats | diatom_chain | plankton |
| Aeshna grandis | dragonflies | Acer platanoides | plants_terrestrial |
| Brachytron pratense | dragonflies | Ajuga reptans | plants_terrestrial |
| Cordulegaster bidentata | dragonflies | Allium ursinum | plants_terrestrial |
| Enallagma cyathigerum | dragonflies | Anemone ranunculoides | plants_terrestrial |
| Erythromma lindenii | dragonflies | Angelica sylvestris | plants_terrestrial |
| Erythromma viridulum | dragonflies | Aruncus dioicus | plants_terrestrial |
| Gomphus vulgatissimus | dragonflies | Barbarea vulgaris | plants_terrestrial |
| Libellula quadrimaculata | dragonflies | Calystegia sepium | plants_terrestrial |
| Onychogomphus forcipatus forcipatus | dragonflies | Cardamine amara | plants_terrestrial |
| Orthetrum brunneum | dragonflies | Carex paniculata | plants_terrestrial |
| Orthetrum cancellatum | dragonflies | Crepis biennis | plants_terrestrial |
| Somatochlora metallica | dragonflies | Dryopteris filix-mas | plants_terrestrial |
| Sympecma fusca | dragonflies | Elymus repens | plants_terrestrial |
| Sympetrum vulgatum | dragonflies | Euonymus europaeus | plants_terrestrial |
| Astacus_astacus | fish | Festuca rubra | plants_terrestrial |
| Carassius_carassius | fish | Frangula alnus | plants_terrestrial |
| Carassius_gibelio | fish | Galium album | plants_terrestrial |
| Cyprinus_carpio | fish | Geranium sylvaticum | plants_terrestrial |
| Gobio_gobio | fish | Helictotrichon pubescens | plants_terrestrial |
| Lepomis_gibbosus | fish | Hieracium murorum | plants_terrestrial |
| Rutilus_rutilus | fish | Lathyrus vernus | plants_terrestrial |
| Alisma plantago-aquatica | macrophytes | Lonicera xylosteum | plants_terrestrial |
| Callitriche cophocarpa | macrophytes | Lysimachia nummularia | plants_terrestrial |
| Carex acuta | macrophytes | Lysimachia vulgaris | plants_terrestrial |
| Carex elata | macrophytes | Phragmites australis | plants_terrestrial |
| Carex riparia | macrophytes | Picea abies | plants_terrestrial |
| Carex vesicaria | macrophytes | Poa angustifolia | plants_terrestrial |
| Ceratophyllum demersum | macrophytes | Polystichum aculeatum | plants_terrestrial |
| Glyceria fluitans | macrophytes | Potentilla verna | plants_terrestrial |
| Lycopus europaeus | macrophytes | Prunus avium | plants_terrestrial |
| Myosotis scorpioides | macrophytes | Rubus armeniacus | plants_terrestrial |
| Poa palustris | macrophytes | Rubus hystrix | plants_terrestrial |
| Polygonum amphibium | macrophytes | Rumex hydrolapathum | plants_terrestrial |
| Potamogeton berchtoldii | macrophytes | Salix viminalis | plants_terrestrial |
| Potamogeton crispus | macrophytes | Sedum sexangulare | plants_terrestrial |
| Ranunculus trichophyllus | macrophytes | Typha latifolia | plants_terrestrial |
| Scrophularia umbrosa | macrophytes | Vicia sepium | plants_terrestrial |
| Sparganium emersum | macrophytes |  |  |
| Utricularia vulgaris | macrophytes |  |  |
| Zannichellia palustris | macrophytes |  |  |
| ciliate_colony | plankton |  |  |
| cladocera | plankton |  |  |
| closterium | plankton |  |  |
| cyclops | plankton |  |  |
| dinobryon | plankton |  |  |
| dinoflagelate | plankton |  |  |
| dinoflagellate_cyst | plankton |  |  |
| gomphonema | plankton |  |  |
| green_algae | plankton |  |  |
| larvae | plankton |  |  |
| nitzschia | plankton |  |  |
| phacus | plankton |  |  |
| Acer campestre | plants_terrestrial |  |  |
| Agrostis gigantea | plants_terrestrial |  |  |
| Alnus glutinosa | plants_terrestrial |  |  |
| Alnus incana | plants_terrestrial |  |  |
| Athyrium filix-femina | plants_terrestrial |  |  |
| Bromus inermis | plants_terrestrial |  |  |
| Bromus sterilis | plants_terrestrial |  |  |
| Cardamine impatiens | plants_terrestrial |  |  |
| Carex brizoides | plants_terrestrial |  |  |
| Carex riparia | plants_terrestrial |  |  |
| Circaea alpina | plants_terrestrial |  |  |
| Circaea lutetiana | plants_terrestrial |  |  |
| Cirsium oleraceum | plants_terrestrial |  |  |
| Clematis vitalba | plants_terrestrial |  |  |
| Convallaria majalis | plants_terrestrial |  |  |
| Deschampsia cespitosa | plants_terrestrial |  |  |
| Epilobium ciliatum | plants_terrestrial |  |  |
| Epilobium montanum | plants_terrestrial |  |  |
| Epilobium parviflorum | plants_terrestrial |  |  |
| Epilobium tetragonum | plants_terrestrial |  |  |
| Equisetum hyemale | plants_terrestrial |  |  |
| Festuca arundinacea | plants_terrestrial |  |  |
| Galium odoratum | plants_terrestrial |  |  |
| Geranium robertianum | plants_terrestrial |  |  |
| Glyceria fluitans | plants_terrestrial |  |  |
| Glyceria notata | plants_terrestrial |  |  |
| Humulus lupulus | plants_terrestrial |  |  |
| Impatiens glandulifera | plants_terrestrial |  |  |
| Impatiens noli-tangere | plants_terrestrial |  |  |
| Juncus effusus | plants_terrestrial |  |  |
| Knautia dipsacifolia | plants_terrestrial |  |  |
| Lapsana communis | plants_terrestrial |  |  |
| Lemna minor | plants_terrestrial |  |  |
| Mentha aquatica | plants_terrestrial |  |  |
| Milium effusum | plants_terrestrial |  |  |
| Poa annua | plants_terrestrial |  |  |
| Polygonum aviculare | plants_terrestrial |  |  |
| Polygonum hydropiper | plants_terrestrial |  |  |
| Potentilla sterilis | plants_terrestrial |  |  |
| Prunus spinosa | plants_terrestrial |  |  |
| Ranunculus repens | plants_terrestrial |  |  |
| Rorippa amphibia | plants_terrestrial |  |  |
| Rosa canina | plants_terrestrial |  |  |
| Rosa multiflora | plants_terrestrial |  |  |
| Rubus idaeus | plants_terrestrial |  |  |
| Salix alba | plants_terrestrial |  |  |
| Salix caprea | plants_terrestrial |  |  |
| Sonchus asper | plants_terrestrial |  |  |
| Sparganium erectum | plants_terrestrial |  |  |
| Stachys palustris | plants_terrestrial |  |  |
| Stachys sylvatica | plants_terrestrial |  |  |
| Torilis japonica | plants_terrestrial |  |  |
| Veronica anagallis-aquatica | plants_terrestrial |  |  |
| Veronica beccabunga | plants_terrestrial |  |  |
| Viburnum lantana | plants_terrestrial |  |  |

*Table S8 Summary of linear model results testing the relationship between human land-use intensity and the nestedness component /total beta diversity, as well as gamma species richness (total species richness per taxa per site). For each taxa the table reports the R² and p-values of the models. Significant relationships (p < 0.05) are marked in bold in the table.*

| **Ratio nestedness/beta diversity** |  |  |  |  |
| --- | --- | --- | --- | --- |
| **community** | **estimate** | **std_error** | **r_squared** | **p_value** |
| arthropods_terrestrial | -5.92E-07 | 1.52E-06 | 0.01 | 0.70 |
| plants_terrestrial | 2.03E-07 | 4.72E-07 | 0.01 | 0.67 |
| aquatic_invertebrates | -2.40E-07 | 6.73E-07 | 0.01 | 0.73 |
| dragonflies | -1.24E-06 | 1.26E-06 | 0.07 | 0.34 |
| fish | -2.23E-06 | 1.76E-06 | 0.19 | 0.25 |
| macrophytes | -4.31E-06 | 1.25E-06 | 0.52 | **< 0.01** |
| plankton | 6.47E-07 | 1.51E-06 | 0.01 | 0.67 |
| arthropods_flying | -2.59E-06 | 2.17E-06 | 0.19 | 0.28 |
| bats | -2.13E-06 | 2.26E-06 | 0.13 | 0.38 |
| **Gamma species richness** |  |  |  |  |
| **community** | **estimate** | **std_error** | **r_squared** | **p_value** |
| amphibians | -1.68E-05 | 4.22E-06 | 0.53 | **< 0.01** |
| aquatic_invertebrates | -2.46E-05 | 4.54E-05 | 0.02 | 0.60 |
| arthropods_flying | 1.08E-05 | 6.28E-06 | 0.33 | 0.14 |
| arthropods_terrestrial | -7.91E-06 | 1.05E-05 | 0.04 | 0.46 |
| bats | -7.01E-06 | 1.17E-05 | 0.06 | 0.57 |
| dragonflies | -2.29E-05 | 2.06E-05 | 0.08 | 0.29 |
| fish | 2.39E-05 | 1.23E-05 | 0.21 | 0.07 |
| macrophytes | -5.41E-06 | 1.64E-05 | 0.01 | 0.75 |
| plankton | -3.41E-05 | 1.96E-05 | 0.18 | 0.10 |
| plants_terrestrial | 4.07E-05 | 3.98E-05 | 0.07 | 0.32 |
